# GENA-LM: A Family of Open-Source Foundational DNA Language Models for Long Sequences

**DOI:** 10.1101/2023.06.12.544594

**Authors:** Veniamin Fishman, Yuri Kuratov, Aleksei Shmelev, Maxim Petrov, Dmitry Penzar, Denis Shepelin, Nikolay Chekanov, Olga Kardymon, Mikhail Burtsev

## Abstract

Recent advancements in genomics, propelled by artificial intelligence, have unlocked unprecedented capabilities in interpreting genomic sequences, mitigating the need for exhaustive experimental analysis of complex, intertwined molecular processes inherent in DNA function. A significant challenge, however, resides in accurately decoding genomic sequences, which inherently involves comprehending rich contextual information dispersed across thousands of nucleotides. To address this need, we introduce GENA-LM, a suite of transformer-based foundational DNA language models capable of handling input lengths up to 36,000 base pairs. Notably, integrating the newly-developed Recurrent Memory mechanism allows these models to process even larger DNA segments. We provide pre-trained versions of GENA-LM, including multispecies and taxon-specific models, demonstrating their capability for fine-tuning and addressing a spectrum of complex biological tasks with modest computational demands. While language models have already achieved significant breakthroughs in protein biology, GENA-LM showcases a similarly promising potential for reshaping the landscape of genomics and multi-omics data analysis. All models are publicly available on GitHub https://github.com/AIRI-Institute/GENA_LM and HuggingFace https://huggingface.co/AIRI-Institute. In addition, we provide a web-service https://dnalm.airi.net/ allowing user-friendly DNA annotation with GENA-LM models.

## 1 Main

The encoding of genetic information by DNA is a principal subject in biology, involving both straightforward and complex systems of translation and epigenetic coding, respectively. While the translation of messenger RNA to amino acid sequences employs a widely-accepted genetic code, other forms of encoding, notably the epigenetic code, are more challenging [1]. DNA sequences dictate functional genome elements, including promoters, enhancers, and transcription factor binding sites, among others. However, the diversity and redundancy of their underlying motifs challenge their detection within vast eukaryotic genomes, complicating insights into non-coding genome evolution and interpretations of human genomic variants, given the yet-to-be-fully-unraveled complexity of the epigenetic code.

The advent of next-generation sequencing and additional high-throughput technologies has catalyzed the accumulation and public deposition of extensive databases, rich with functional genomic elements, enabling the broad application of computational methods to large-scale genomic data analysis [2]. We, along with others [3], have successfully employed machine-learning methods, including ensemble learning [4] and convolutional neural networks [5, 6], for this purpose. However, while potent, these approaches encounter constraints in identifying long-range dependencies within DNA sequences, a common phenomenon in human and other eukaryotic genomes [7]. Recent strategies employing transformer neural network-based approaches seek to surmount these constraints [8], with cutting-edge transformer architectures showcasing the capability to infer specific epigenetic properties and gene expression levels from DNA sequences with exceptional precision [8]. Nonetheless, a primary limitation exists in that training these specialized in domain models requires significant computational resources, and their inference capabilities are bounded by the features integrated into the training dataset.

Transfer learning, especially through pre-training, has been widely adopted in natural language processing for its capacity to enhance computational efficiency and performance in scenarios with limited target data [9–14]. Models pre-trained on substantial unlabeled datasets can be fine-tuned or utilized as feature extractors for new tasks, frequently outperforming models trained on task-specific datasets, particularly when those datasets are smaller. The application of this approach to bioinformatics is exemplified by the development of DNABERT [15], a BERT-like transformer neural network [14, 16] pre-trained on the human genome to predict subsequences from context, and subsequently fine-tuned for downstream tasks such as promoter activity prediction and transcription factor binding. While DNABERT signifies a promising advance, its applicability is hindered by an input size cap of 500 base pairs, and recent DNABERT extension DNABERT-2 [17] also has an input length limit of approximately 1-4 kb. This input length limitation restricts the ability of the models to capture the extended contexts vital for various genomic applications.

Enhancing input size for transformer models has recently been addressed through several developments, including sparse attention, effective attention, and recurrence. Sparse attention techniques, which utilize either predefined or learned attention patterns like sliding window or block-diagonal, linearize the quadratic dependency of full attention on input length [18–22]. Conversely, linear attention methods approximate full token-to-token interactions through softmax linearization [23, 24]. In the domain of recurrent models, inputs are segmented and sequentially processed, with inter-segment information relayed through prior hidden states [25, 26] or specialized memory [27–29]. Notably, the recently introduced Recurrent Memory Transformer architecture facilitates information aggregation from both long [29] and extremely long input sequences [30], spanning thousands to millions of elements respectively.

In this work, we introduce GENA-LM, a family of transformer-based foundational DNA models. After fine-tuning for predictive analysis of various functional genomic elements — including promoter activity, splicing, polyadenylation sites, enhancer annotations, and chromatin profiles — GENA-LM models demonstrate state-of-the-art performance for a significant fraction of tasks, achieving top average performance relative to other models. Moreover, our augmentation of GENA-LM with the Recurrent Memory Transformer (RMT) enables tackling genomic tasks that require substantial input sequence lengths. We also explore new applications of GENA-LM, such as identifying DNA motifs essential for transcription factor binding and assessing mutation effects in promoters and splice sites to aid in the prioritization of clinical variants. To broaden the model’s utility beyond human genomes, we have developed and released species-specific models for yeast, Arabidopsis, and Drosophila, as well as a multispecies model. To facilitate sequence annotation for the thousands of unannotated sequences now available, we have developed GENA-Web^1^, a web service that generates various annotations based on DNA sequence input. We contribute to the research community by open-sourcing the GENA-LM family on GitHub^2^ and providing pre-trained models (prefixed with gena-lm-) on Hugging Face^3^.

## 2 Results

### 2.1 Family of pre-trained transformer based GENA-LM models

In this study, we introduce a new universal transformer model tailored for nucleic acid sequences, which offers several improvements over existing models such as DNABERT [15] and BigBird [21] (as depicted in Fig. 1, A). To ensure its versatility across various applications, we pre-trained our model using multiple datasets and diverse input sequence lengths.

**Fig. 1.**
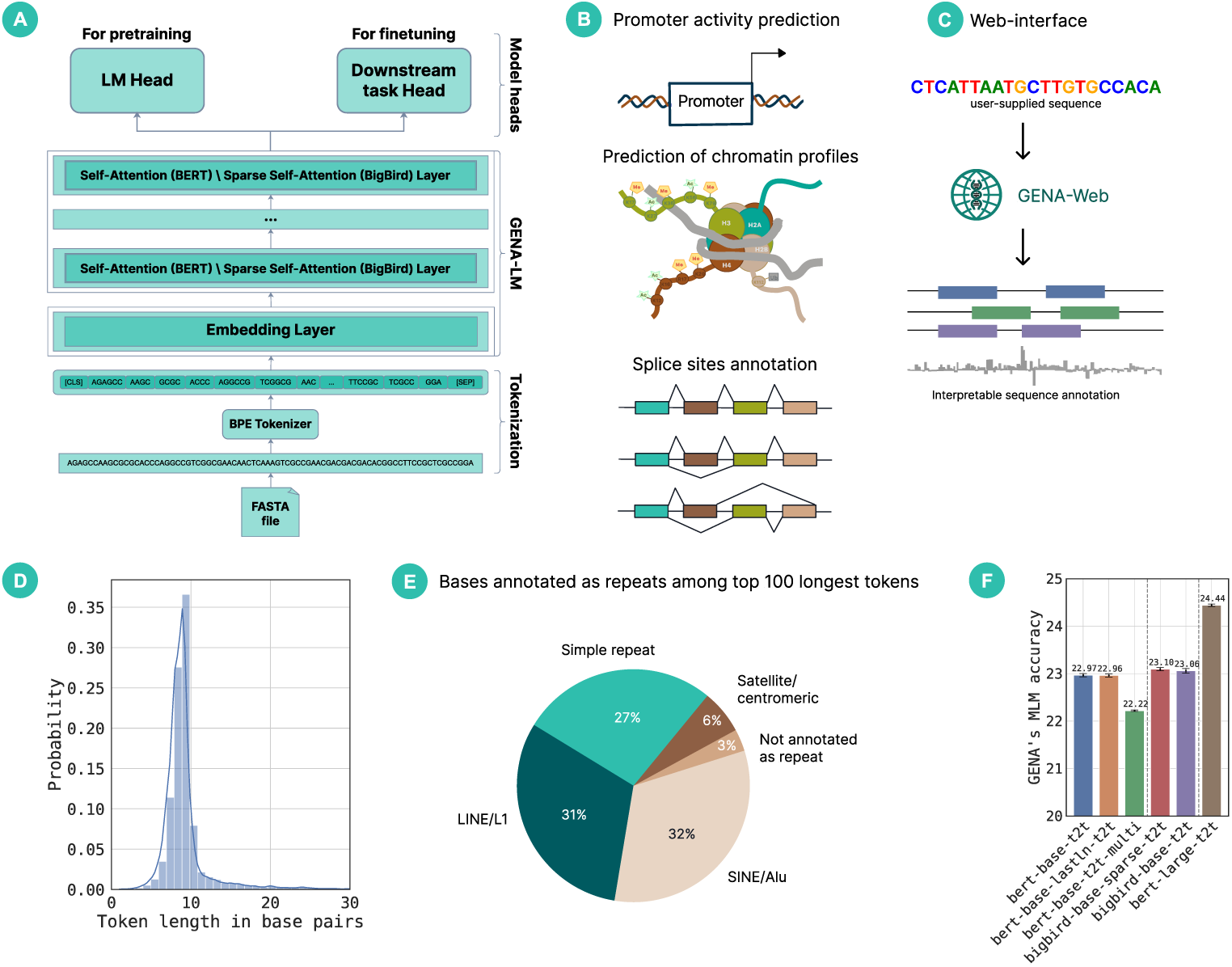
The GENA-LM Family of Foundational DNA Language Models. **A**. The GENA-LM transformer-based architecture is pre-trained on DNA sequences using a masked language modeling (MLM) objective. GENA-LMs encompass a variety of models that differ in their pre-training data and architecture, as detailed in Table 4. All models adhere to the same workflow: DNA sequences are tokenized using a BPE algorithm before being processed through transformer layers, which generate representations of the input sequences that are suitable for downstream applications. Post pre-training, this foundational DNA model incorporates a downstream task-specific head, which utilizes DNA representations to address specific genomic tasks during the fine-tuning process. **B**. GENA’s evaluation tasks include predictions related to promoter and enhancer activities, splicing sites, chromatin profiles, and polyadenylation site strength (not all shown). **C**. Task specific fine-tuned models can be queried via web service^1^. **D**. Post-BPE tokenization, the median token length stands at nine base pairs, as reflected in the token length distribution. **E**. Illustration of repetitive element representation for the 100 longest tokens. **F**. GENA’s model accuracies for pre-training on the masked language modeling task demonstrate that models with a higher parameter count achieve superior performance.

In the data preprocessing phase, we have extended established pipelines by integrating Byte-Pair Encoding (BPE) for sequence tokenization (Fig.1, A, bottom). The essence of BPE is that it constructs a sequence dictionary to pinpoint the most frequently occurring subsequences within the genome. This results in tokens of diverse lengths, ranging from a single base pair up to 64 base pairs. In our tests, the median token length was determined to be 9 base pairs (Fig.1, D). Interestingly, our BPE vocabulary revealed tokens of significant biological relevance. For example, the longest tokens were often indicative of familiar repetitive elements, such as LINEs or simple repeats (Fig. 1, E). The tokenization approach we adopted is greedy, starting with the longest sequences in the dictionary and tokenizing them first. Employing non-overlapping tokens, as opposed to the overlapping k-mers used in earlier studies, allows for the analysis of more extended sequence fragments while maintaining the same model input size. To put it in perspective, 512 overlapping 6-mers represent 512 base pairs, but 512 non-overlapping BPE tokens can represent approximately 4.5 kb. This is a crucial factor when dealing with expansive and intricate genomes like that of humans. Nevertheless, it’s worth noting that the model’s granularity is confined to the resolution of these individual tokens, which might pose constraints for certain applications.

Our second enhancement pertains to the diversification in the implementation of the attention mechanism. The foundational GENA models utilize a conventional attention mechanism, which empowers the model to discern relationships between every pair of tokens in the input sequences. Conversely, Sparse GENA models incorporate a sparse attention mechanism. This approach extends the permissible length of the input sequence by constraining the overall number of connections. Nevertheless, it retains the capability to understand relationships between distant sequence elements. In the case of recurrent GENA models (Section 2.6), the transformer is supplemented with memory capabilities. This modification facilitates the processing of even longer inputs by segmenting them.

Through the integration of BPE tokenization and the sparse attention mechanism, we are able to train models that can handle input sequences of approximately 4.5 kb (512 tokens with full attention) and 36 kb (4096 tokens with sparse attention). The incorporation of recurrent memory further expands this capacity, allowing for the processing of input sequences spanning hundreds of thousands of base pairs.

For model training, we utilized the masked language modeling task, a prevalent technique in natural language processing wherein the model predicts a masked token based on its surrounding sequence context. Unlike previous studies that used the hg38 genome assembly [15, 21], we trained all our models using the more recent human T2T genome assembly, setting our experiment apart. To mitigate the risk of overfitting to the reference genome, we incorporated common variants from the 1000-genome project database into some of our models. Additionally, we enriched our training dataset with genomes from diverse species, encompassing standard model organisms such as mice, fruit flies, nematode worms, and baker’s yeast, as well as others covering the entire spectrum of eukaryotic taxa. For a detailed methodology see Section 4.

Throughout the manuscript, we collectively refer to our suite of developed models as GENA language models (GENA-LMs). Each specific model is designated by its label as shown in Table 4. While each model has its unique merits and constraints, we wish to highlight the following:

1. The *gena-lm-bert-base-t2t* model: This model emulates the BERT transformer architecture, serving as a benchmark for subsequent models.
2. The *gena-lm-bert-base-t2t-yeast/fly/athaliana/multi* models: These models include multispecies or taxon-specific data during pre-training while using the same BERT architecture as a model described above.
3. The *gena-lm-bert-large-t2t* model: With the most significant parameter count (336M) and an input capacity of 4.5 kb, it stands out in terms of complexity.
4. The *gena-lm-bigbird-base-sparse-t2t* models: These models, although having fewer parameters than the *gena-lm-bert-large-t2t*, boast a more extended input sequence length of 36 kb.

Upon evaluating the performance of our models in the masked language modeling task (Fig. 1, F), we observed that models with sparse attention slightly outperformed their full-attention counterparts limited to 512 tokens. This underscores the role of contextual information in the training regimen. Nonetheless, it’s imperative to note that while achieving commendable scores in the masked language modeling task is encouraging, it doesn’t necessarily guarantee optimal translation of the learned DNA representations to downstream applications. Consequently, our study delves into the comprehensive assessment of GENA-LMs across a spectrum of biologically relevant tasks to explore their merits and constraints.

### 2.2 GENA-LM performance on different genomic tasks

To evaluate the foundational GENA-LM models, we selected a range of genomic challenges that have recently been addressed using artificial intelligence (Fig. 1, B). These challenges encompass 1) prediction of polyadenylation site strength; 2) Forecasting of chromatin profiles, which includes histone modifications, DNase I hypersensitivity sites and transcription factor binding sites, among others; 3) Identification of splicing sites. In addition, we employed a comprehensive set of 18 benchmarks recently developed by [31]. The datasets for these tasks are derived from human genomic data. To explore the performance of models when applied to non-vertebrate species, we introduced challenges including 4) determining the activity of housekeeping and developmental enhancers in a STARR-seq assay in *Drosophila* cells [32] and 5) estimation of DNA sequence promoter activity in humans, flies, yeasts, and plants.

#### Prediction of chromatin profiles

The major feature of GENA-LMs is their ability to process long DNA sequences, ranging from 4 to 36 kb. Consequently, we benchmarked GENA-LMs on tasks where understanding longrange dependencies in DNA sequences is crucial for accurate prediction. In genomics, these long-range dependencies are particularly significant for various epigenetic features, which makes predicting a locus’s epigenetic states based on its sequence a significant challenge. To assess the capabilities of the GENA-LM transformers in addressing this, we used DeepSEA dataset[33]. This dataset encompasses over 900 cell-type-specific chromatin profiles, which are grouped into DNAse I hypersensitivity sites (DHS), histone marks (HM), and transcription factor binding sites (TF). In the foundational DeepSEA challenge, chromatin mark signals were predicted for each 200 bp genomic segment, informed by both its sequence and an additional 800 bp context derived from its flanking regions (±400 bp).

When deploying GENA-LMs for this challenge (see DeepSEA section of Table 1), we discovered that transformer models markedly surpassed the performance metrics previously achieved by the convolutional neural network, DeepSEA. Notably, for TF and DHS profiles, GENA-LMs delivered scores that eclipsed those reported for the BigBird architecture, even though Big-Bird utilized an expanded 8 kb context (leading GENA-LM average AUC on a 1 kb context for TF: 96.81 ± 0.1 vs. BigBird’s 96.1; for DHS: 92.8 ± 0.03 vs. BigBird’s 92.3). Furthermore, the performance metrics for GENA-LMs were either on par with or exceeded those recently reported for the Nucleotide Transformer [31]. They also proved superior to the outcomes of the DNABERT architecture when trained on 1 kb input lengths.

**Table 1.** Comparative performance analysis of GENA-LM models across multiple genomic tasks and input sizes. This table encapsulates results for tasks including: DeepSEA [33] chromatin profile prediction (DNase I hypersensitivity (DHS), histone modifications (HM), and transcription factor (TF) binding sites); Promoter activity prediction based on EPDnew dataset; and splice sites annotation based on SpliceAI dataset. All values are average of at least three runs.

| Sub-task \ input size | GENA-LM models |  |  |  |  |  | DNABERT Models |  |
| --- | --- | --- | --- | --- | --- | --- | --- | --- |
|  | bert-base-t2t | bert-base-lastln-t2t | bert-base-multi-t2t | bert-large-t2t | bigbird-base-t2t | bigbird-base-sparse-t2t | DNABERT | DNABERT-2 |
| DeepSEA chromatin profiling, ROC AUC |  |  |  |  |  |  |  |  |
| DHS \ 1kb |  | 92.15 | 89.13 | **92.87** | 90.73 | 92.04 | 92.70 | 86.03 |
| DHS \ 8kb | - | - | - | 87.85 | 92.26 | 92.06 | - | - |
| HM \ 1kb | 85.17 | 86.05 | 82.65 | 86.64 | 84.72 | 85.31 | 86.13 | 79.14 |
| HM \ 8kb | - | - | - | 88.18 | **89.71** | 89.69 | - | - |
| TF \ 1kb | 95.69 | 95.98 | 92.83 | 96.54 | 94.96 | **96.81** | 96.37 | 88.53 |
| TF \ 8kb | - | - | - | 95.18 | 96.24 | 96.40 | - | - |
| EPDnew promoter activity, f1 |  |  |  |  |  |  |  |  |
| promoter \ 0.3 kb | 89.88 | 90.08 | 89.70 | 90.62 | 89.98 | 87.57 | 93.26 | 89.44 |
| promoter \ 2 kb | 93.41 | 93.62 | 93.11 | 94.16 | 93.42 | 93.85 | **94.28** | 92.98 |
| promoter \ 16 kb | - | - | - | - | 93.15 | 93.40 | - | - |
| Splice site annotation, AP score |  |  |  |  |  |  |  |  |
| splice site \ 15 kb | 92.63 | 92.56 | 91.42 | 93.6 | **94.78** | 94.7 | - | - |

To ensure a more equitable comparison between the BigBird and GENA-LM architectures, we adapted the DeepSEA dataset to incorporate expanded context sequences. This adaptation allowed us to match the 8 kb input length characteristic of the BigBird architecture. Intriguingly, the augmented context had differential effects on the prediction of various epigenetic profiles. For histone marks, a marked performance improvement was evident, with an AUC reaching 89.71 ± 0.08. This was notably superior to the shorter context’s score of 86.64 ± 0.08, the original DeepSEA findings (85.6), and the BigBird’s result (88.70). However, for TF and DHS predictions, the extension in input length yielded only marginal enhancements in performance.

We analyzed the AUC variations across individual histone marks to pinpoint which epigenetic profiles were responsible for the observed performance enhancement. Remarkably, there was a distinct divergence between narrow and broad histone marks. While the narrow marks demonstrated marginal AUC improvements, the broad marks exhibited a pronounced increase when the context length was extended (Supplementary Fig. 1).These observations reinforce our prior observation [5] that broad histone marks necessitate an expansive context for precise prediction. This highlights the importance of handling extended input lengths for such tasks. The varying performance metrics of distinct GENA-LMs across diverse epigenetic profiles and context lengths underscore that no singular model universally excels across all challenges. For transcription factors (TFs), the *gena-lm-bigbird-base-sparse-t2t* stands out on 1 kb inputs, with performance diminishing marginally when the input size increases. In contrast, for DHS, the *gena-lm-bert-large-t2t* model, boasting the highest parameter count, emerges as the optimal choice. Surprisingly, extending the context for this model results in a notable performance dip. For histone marks (HM), the optimal approach hinges on processing extended contexts with the *gena-lm-bigbird-base-t2t* model. Collectively, GENA-LMs outstrips competing models like BigBird, DNABERT, and Nucleotide Transformer, marking a new performance state of the art for this task.

#### Promoter activity prediction

Promoter activity is an essential characteristic of genomic sequences, allowing them to drive the expression of genes. Although the basal promoter is a relatively short genomic region of °300 bp., surrounding context can substantially modify promoter activity. We assessed whether our models can discriminate human promoter sequences using promoter instances from the EPD dataset and juxtaposed them against non-promoter control samples. We observed that when the input sequence length was extended from 300 bp to 2 kb, there was a significant improvement in performance, as shown in the EPDnew section of Table 1. With 300 bp sequences, the DNABERT architecture emerged superior, registering an f1 score of 93.26, compared to the top-performing GENA-LM’s f1 score of 90.62. However, when evaluating 2 kb sequences, the GENA-LM performance matched DNABERT results, recording an f1 score of 94.28 ± 0.65 and 94.16 ± 0.19 for DNABERT and *gena-lm-bert-large-t2t*, respectively (no significant difference, Wilcoxon test p-value=0.0625). This result was markedly higher than the DNABERT-2 model’s score, which, when fine-tuned for the same input sequence length, achieved an f1 score of 92.98 ± 0.25.

In assessing the performance of GENA-LMs for predicting promoter activity, we observed the following: 1) Models with a greater number of parameters outperformed those with fewer. For instance, the *gena-lm-bert-large-t2t* surpassed the *gena-lm-bert-base-t2t*. 2) The ability to handle longer input sequences due to the sparse attention mechanism gave certain models an edge over traditional full-attention BERT models. As a result, the *gena-lm-bigbird-base-sparse* outperformed the *gena-lm-bert-base-t2t*. Interestingly, models with shorter inputs but more parameters, such as the *gena-lm-bert-large-t2t*, still had superior performance over the *gena-lmbigbird-base-sparse*. 3) Incorporating multispecies training by using genomic sequences beyond just human data during pre-training did not result in improved performance, as seen when comparing the *gena-lm-bert-base-t2t-multi* with the *gena-lm-bert-base-t2t*.

#### Splice site annotation

We further optimized GENA-LMs to predict splice-donor and splice-acceptor sites within the human genome (Splice section of Table 1). The task required analyzing large contexts: a 15 kb input comprised of a central 5 kb target flanked by 5 kb sequences on either end. Notably, the task-specific convolutional neural network, SpliceAI, marginally surpassed GENA-LMs, registering a mean PR AUC of 0.960 compared to 0.947 ± 0.002 for GENA-LMs.

For this task, models designed for longer sequence inputs, such as *gena-lm-bigbird-base-t2t*, outperformed those tailored for shorter inputs, even if the latter were equipped with more parameters, as in *gena-lm-bert-large-t2t*. This aligns with our earlier findings, suggesting that extending contextual information could be more beneficial than merely increasing the number of parameters. Consistent with our promoter analysis, multispecies models, like *gena-lm-bertbase-t2t-multi* (mean PR AUC of 0.914), did not enhance performance compared to their single-species counterparts, such as *gena-lm-bert-base-t2t* (mean PR AUC of 0.926).

#### Benchmarking GENA-LMs on short sequence tasks

To compare GENA-LM with with several recently developed DNA language models, including Nucleotide Transformer [31], DNABERT [15], DNABERT-2 [17], HyenaDNA [34], and finetuned versions of Enformer [8] we adopted a recent series of 18 benchmarks [31], which include relatively short sequence inputs ranging from 300 to 600 bp. According to the results presented in the Table 2, *gena-lm-bert-large-t2t* outperformed all other models, achieving the highest average score and the second-best average ranked score. Notably, *gena-lm-bert-large-t2t* outperformed Nucletide Transformer in multiple tasks, despite having substantially less parameters (330M vs 2500M). Similarly, GENA-LMs demonstrated performance *on par* with these models in another series of benchmarks focused on prediction of the human polyadenylation sites and *Drosophila* enhancer activity, as detailed in Supplementary Note 1.

**Table 2.** GENA-LM (*gena-lm-bert-large-t2t* ) has the highest average score over the wide range of genomic tasks compared to other foundational DNA models. Datasets and performance of alternative models for 18 tasks are form [31]. All values are MCCs.

| Task \ Dataset | GENA- bert-<br>large-t2t | Model |  |  |  |  |  |  |  |  |  |  |  |  |
| --- | --- | --- | --- | --- | --- | --- | --- | --- | --- | --- | --- | --- | --- | --- |
|  |  | HyenaDNA-1Kb | HyenaDNA-30Kb | DNABERT-1 | DNABERT-2 | Enformer | NT-HumanRef (500M) | NT-1000G (500M) | NT-1000G (2.5B) | NT-Multispec (2.5B) | NT-Multispec-v2 (500M) | NT-Multispec-v2 (250M) | NT-Multispec-v2 (100M) | NT-Multispec-v2 (50M) |
| H3 | **0.79** | 0.78 | 0.74 | 0.76 | 0.79 | 0.72 | 0.72 | 0.74 | 0.76 | **0.79** | 0.78 | **0.79** | **0.79** | **0.79** |
| H3K14ac | 0.60 | **0.61** | 0.4 | 0.4 | 0.52 | 0.29 | 0.37 | 0.38 | 0.45 | 0.54 | 0.55 | 0.54 | 0.52 | 0.51 |
| H3K36me3 | 0.61 | 0.61 | 0.48 | 0.47 | 0.59 | 0.34 | 0.45 | 0.47 | 0.53 | 0.62 | **0.63** | 0.62 | 0.59 | 0.58 |
| H3K4me1 | 0.53 | 0.51 | 0.38 | 0.4 | 0.51 | 0.29 | 0.37 | 0.38 | 0.42 | 0.54 | **0.55** | 0.54 | 0.52 | 0.52 |
| H3K4me2 | **0.46** | **0.46** | 0.28 | 0.28 | 0.34 | 0.21 | 0.26 | 0.26 | 0.28 | 0.32 | 0.32 | 0.32 | 0.3 | 0.3 |
| H3K4me3 | **0.55** | **0.55** | 0.29 | 0.26 | 0.35 | 0.16 | 0.24 | 0.24 | 0.31 | 0.41 | 0.41 | 0.4 | 0.38 | 0.33 |
| H3K79me3 | **0.67** | **0.67** | 0.57 | 0.58 | 0.61 | 0.5 | 0.56 | 0.56 | 0.57 | 0.62 | 0.63 | 0.62 | 0.61 | 0.6 |
| H3K9ac | **0.61** | 0.58 | 0.47 | 0.5 | 0.54 | 0.42 | 0.45 | 0.46 | 0.49 | 0.55 | 0.56 | 0.55 | 0.54 | 0.52 |
| H4 | 0.78 | 0.76 | 0.76 | 0.79 | 0.8 | 0.73 | 0.75 | 0.76 | 0.79 | **0.81** | 0.8 | 0.8 | 0.79 | 0.8 |
| H4ac | **0.59** | 0.56 | 0.38 | 0.36 | 0.46 | 0.27 | 0.33 | 0.34 | 0.41 | 0.49 | 0.5 | 0.5 | 0.48 | 0.46 |
| Enhancers | **0.55** | 0.52 | 0.49 | 0.5 | 0.52 | 0.45 | 0.5 | 0.51 | 0.54 | 0.55 | 0.55 | 0.54 | 0.5 | 0.52 |
| Enhancers (types) | **0.45** | 0.39 | 0.36 | 0.37 | 0.42 | 0.31 | 0.43 | 0.4 | 0.44 | 0.45 | 0.42 | 0.45 | 0.41 | 0.4 |
| Promoters | 0.94 | 0.92 | 0.91 | 0.92 | 0.94 | 0.91 | 0.9 | 0.9 | 0.93 | **0.95** | **0.95** | **0.95** | 0.94 | 0.92 |
| Promoters (TATA) | 0.91 | 0.88 | 0.87 | 0.91 | 0.91 | 0.92 | 0.89 | **0.87** | 0.91 | 0.92 | **0.93** | 0.92 | 0.91 | 0.89 |
| Promoters (non-TATA) | 0.94 | 0.92 | 0.91 | 0.92 | 0.94 | 0.91 | 0.91 | 0.9 | 0.93 | **0.95** | **0.95** | **0.95** | 0.94 | 0.92 |
| Splicing (Both) | 0.91 | 0.94 | 0.94 | 0.96 | 0.91 | 0.77 | 0.96 | 0.95 | 0.96 | **0.97** | **0.97** | **0.97** | **0.97** | 0.96 |
| Splicing (Acceptors) | 0.92 | 0.92 | 0.92 | - | 0.95 | 0.83 | 0.93 | 0.93 | 0.96 | **0.97** | 0.96 | 0.96 | 0.96 | 0.95 |
| Splicing (Donors) | 0.91 | 0.9 | 0.92 | - | 0.93 | 0.81 | 0.94 | 0.94 | 0.96 | **0.97** | 0.97 | 0.97 | 0.96 | 0.94 |
| Average | **0.707** | 0.693 | 0.615 | 0.586 | 0.668 | 0.547 | 0.609 | 0.612 | 0.647 | 0.690 | 0.691 | 0.688 | 0.673 | 0.662 |

### 2.3 Identifying functional genomic elements with GENA-LMs

#### Spotting motifs for transcription factors binding

Modern techniques for analyzing deep neural networks allow us to assess the contribution of each input element to a model’s downstream task performance. Such analyses offer valuable insights into the underlying mechanisms of biological processes. Take the ChIP-seq technique, for instance, a prevalent method for chromatin profiling. Its resolution is approximately 100– 200 bp. However, recognition motifs for the majority of DNA-binding proteins are significantly shorter, typically between 4–10 bp. Consequently, deducing precise binding locations from ChIP-seq data is challenging and often necessitates supplementary experiments [35].

To ascertain if GENA can enhance the resolution of experimental ChIP-seq data, we employed token importance scoring [36] on the *bigbird-base-sparse-t2t* model, which was fine-tuned using the DeepSEA dataset. This methodology assigns a significance value to each token within the input, based on its relevance to the prediction outcome. Here, we concentrate on the binding of three transcription factors: ATF1, CTCF, and GATA2 in human K562 cells. Each of these factors possesses well-established DNA recognition motifs (Fig.2, A). This allows for a comparison between tokens important for GENA’s predictions and the recognized sequence determinants associated with transcription factor binding.

**Fig. 2.**
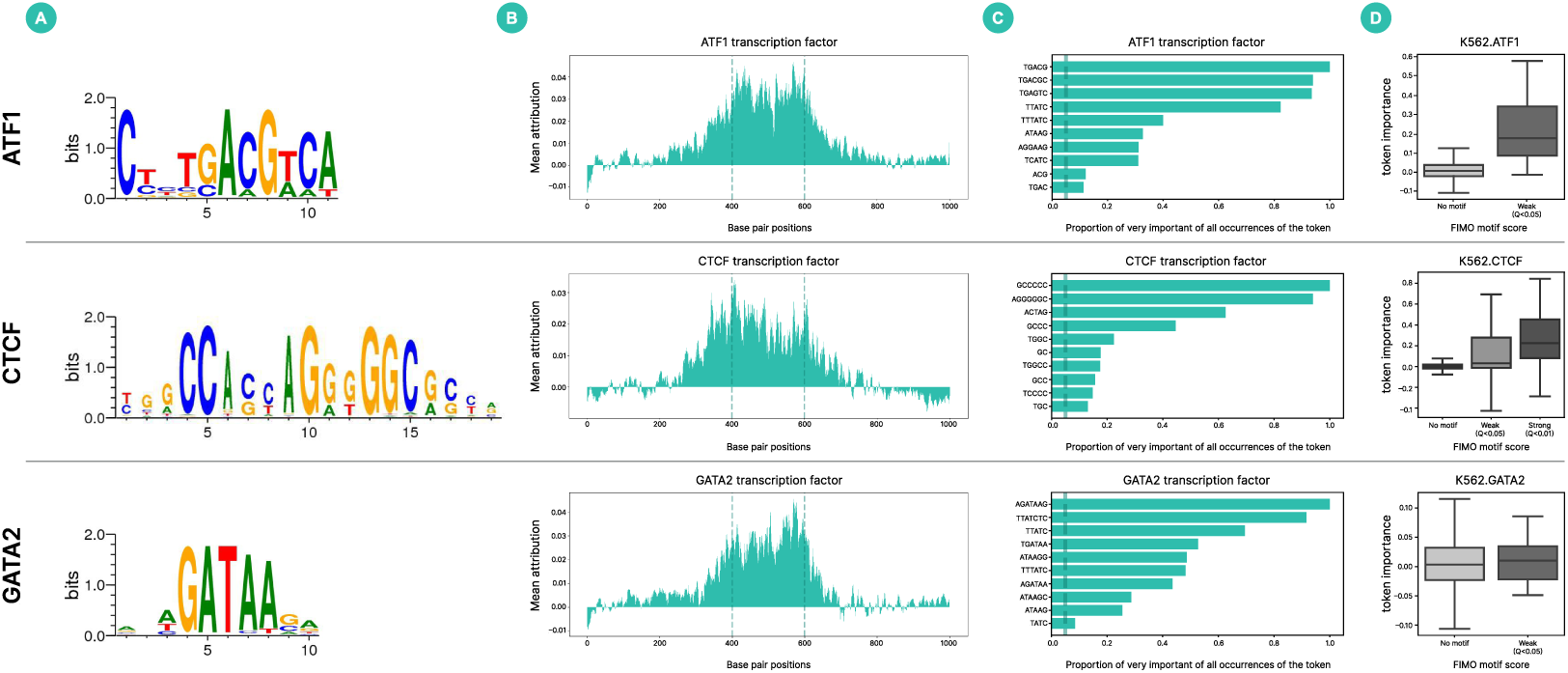
GENA-LM identifies DNA motifs essential for transcription factor binding. In panels A, B, C, and D, each row pertains to a distinct factor, labeled to the left. **A**. Logo representation of motifs for the three transcription factors considered in our analysis. **B**. Profile of average token importance scores over the sequence length. Vertical dashed lines demarcate the 200 bp prediction region. **C**. Bars represent the frequency of token occurrences in the ”highly important” category (tokens with scores in the top 5th percentile). The X-axis shows the proportion of these occurrences relative to all occurrences for that token. A vertical reference line marks the 0.05 fraction threshold; only tokens exceeding this fraction are displayed. **D**. Boxplots detail the distribution of importance scores for tokens, categorized by different FIMO q-values. They display the median, interquartile range, as well as the 5th and 95th percentiles.

First, we examined the distribution of importance scores across the sequence length, as depicted in Fig.2, B. It should be noted that during the fine-tuning process, the input consists of the DNA sequence from the 200 bp target region (where transcription factor binding is anticipated) accompanied by an 800 bp contextual sequence. Our analysis, presented in Fig.2, B, reveals a consistent pattern: the importance attributed to a token diminishes as its distance from the target region increases, a trend observed for all three transcription factors.

We subsequently sought to discern which sequences garnered high token importance scores. Tokens exceeding the 95th percentile of the importance score distribution were designated as ”highly important.” We then compiled tokens that consistently featured on this ”highly important” list. Upon visual examination (Fig. 2, C), we observed that these tokens frequently encompassed full or fragmented motifs of the target transcription factors. For instance, ATF1, which has a core motif of TGACG, prominently displayed a token matching this exact sequence among its highly important tokens. In the case of the GATA2 factor (core motif: GATAA), the token AGATAAG, incorporating the GATA2 motif, was most prevalent among the highly important tokens. As for CTCF, which boasts a motif more intricate and extended than its counterparts, the most recurrent highly important tokens primarily featured GC-rich subsequences of the motif.

To more comprehensively assess the congruence between known transcription factor motifs and ”highly important” tokens, we employed the FIMO tool to annotate all DNA samples. FIMO is a bioinformatics software designed to identify specific motifs by leveraging the motif’s position weight matrix (PWM). As depicted in Supplementary Fig. 2, there’s a discernible overlap between significant tokens and motifs detected by FIMO. Our statistical evaluation establishes a relationship between FIMO motif scores and token importance scores. Both robust motifs (FIMO q-value *<* 0.01) and more tenuous motifs (0.01 *<* FIMO q-value *<* 0.05) manifest markedly elevated token importance scores compared to tokens absent of any discerned motif (FIMO q-value *>* 0.01) (Fig. 2, D). It’s worth noting, in the context of the GATA2 transcription factor which is characterized by a shorter motif length, sequences with high FIMO q-values are absent. Nonetheless, we observed that the majority of the ”highly important” tokens encompass the core GATA2 motif, as delineated in Supplementary Fig. 3.

While the results affirm that token importance mirrors the presence of established motifs for DNA-binding transcription factors, the congruence between FIMO-detected motifs and tokens with high scores isn’t absolute. This observation prompted us to delve into the nature of motifs encompassed by tokens vital for GENA model prediction, yet devoid of the target transcription factor’s annotated motif as per FIMO. Utilizing the *de novo* motif discovery tool XSTREME, we analyzed a subset of important tokens lacking a canonical motif (with FIMO target factor motif q-value *>* 0.05) and examined the enriched motifs therein. Intriguingly, for both CTCF and GATA2, XSTREME predominantly identified their respective motifs. In the case of important ATF1 tokens sans ATF1 motif, the secondary most abundant motif discerned belonged to the ATF-family. This suggests that the rudimentary position weight matrix statistics employed by FIMO might overlook biologically pertinent motif variants that diverge notably from the consensus represented by the PWM. Conversely, GENA-LM exhibits a promising potential in recognizing these variant motifs. When consolidated during XSTREME analysis, these diverse motif representations converge to echo the canonical motif’s PWM. Moreover, our analysis revealed a significant presence of GATA2 motifs within tokens deemed essential for the ATF1 factor, hinting at a possible functional synergy between these transcription factors in K562 cells — a nuance discerned by GENA-LM. Given that ATF1 is an integral part of the AP-1 complex, our findings resonate with, and potentially elucidate, prior experimental data evidencing cooperation between GATA2 and the AP-1 complex [37].

#### Searching for DNA sequence determinants of hisone modifications

Having ascertained GENA’s capacity at pinpointing DNA motifs crucial for transcription factor binding, we turned our attention to discerning sequence determinants associated with histone marks, which currently lack identifiable binding motifs. Our focus centered on H3K4me1, H3K9me3, and H3K27me3. These factors represent histone modifications with distinct and well-documented functional implications: H3K4me1 delineates active genomic regions, H3K9me3 signifies heterochromatin, and H3K27me3 demarcates the suppressed “facultative heterochromatin” including genes under developmental regulation, bound by polycomb group proteins. In our examination of these factors, we evaluated the distribution of importance scores relative to sequence length and accumulated tokens that frequently exhibited high importance values.

As depicted in Supplementary Fig. 4, the distribution of token importance scores across sequence lengths for these epigenetic markers mirrors that of the previously discussed factors. Notably, specific tokens consistently emerged as highly significant in predicting these histone marks, reminiscent of the motif-rich tokens previously identified as important for predicting associated transcription factors (see Supplementary Fig. 5).

Prompted by our observations, we sought to determine if motifs enriched among the tokens with high importance scores were shared across these three factors. Extending the sequences of our selected highly important tokens by 4 bp, we then undertook a rigorous motif analysis using XSTREME. Our examination revealed several motifs of significance (see Supplementary Table 1), each aligning with known transcription factors. For the active promoter mark, H3K4me1, the significant tokens were found to encompass motifs corresponding to the GATA, JUN, and FOSL transcription factors. These findings align with the documented roles of these factors in regulating transcription and influencing the oncogenic transformation of K562 cells [38, 39]. In the context of the H3K27me3 mark, which signifies facultative heterochromatin and designates functional elements repressed in specific cell lineages, our data from hematopoietic K562 cells indicated an enrichment of transcription factor motifs not typically associated with blood cells. Examples include SNAI2, pivotal in epidermal cell differentiation, and ASCLI, a critical regulator of neurogenesis. This suggests that within this setting, GENA discerned genomic motifs that might be activated in alternative cell types but are designated for repression in the K562 lineage. Lastly, for the H3K9me3 mark indicative of constitutive heterochromatin, our analysis highlighted an enrichment of the ZNF274 transcription factor motif. This is in agreement with its established role as a transcriptional repressor.

In summary, our findings indicate that beyond predicting the epigenetic profiles of a specific locus, GENA models can effectively identify the distinct subsequences that drive observed epigenetic signals. Such analysis holds potential to substantially augment the resolution of prevailing experimental approaches, like ChIP-seq, and to pinpoint transcription factors linked to particular histone modifications.

#### Evaluation of clinical relevance of mutations

The capability of GENA-LMs to accurately predict promoter activity inspired us to investigate whether model predictions could functionally characterize variants in human promoters. We compiled ClinVar mutations that overlap with promoter sequences and evaluated them using the odds ratio of promoter presence probability for wild-type versus mutated sequences. While not all pathogenic ClinVar variants overlapping promoters impact predicted promoter activity, a pathogenic-versus-benign classifier based on *gena-lm-bert-large-t2t* predictions achieved an AUC of 0.66 and an average precision of 0.59 (Supplementary Fig. 6). Further, we applied the integrated gradients method [36] to identify input sequences tokens that are the most important for predicting promoter presence. Our analysis revealed that pathogenic mutations are enriched approximately 2.5 times in the top percentile of the most important tokens (p-value *<* 1e-15, Supplementary Table 2). Comparable results were obtained with the *gena-lm-bigbird-base-sparse-t2t* model.

Similar to promoters, mutations at splice sites often have clinical implications in human diseases. To determine whether GENA-LMs can detect the impact of single-nucleotide perturbations at splice donor and acceptor sites, we conducted comprehensive *in silico* mutagenesis, evaluating the effects of every possible single nucleotide substitution within a ± 20 bp sequence surrounding splice sites. Despite the token-level resolution of the inputs, predictions from *genalm-bigbird-base-t2t* proved sensitive to single-nucleotide substitutions. We observed that the model distinctly identifies substitutions within canonical splice site motifs from other single-nucleotide variants (Supplementary Fig. 7). While the latter rarely alter the model’s prediction, mutations within canonical splice sites almost invariably abolish the predicted acceptor or donor site. Notably, the predicted effects of single nucleotide substitutions align precisely with known splice site motifs: changes in the constant motif positions have more significant effects compared to the substitutions in the more variable regions of the motif. These findings highlight the potential of GENA-LMs for clinical interpretation of human genetic variants.

### 2.4 GENA-LMs beyond human species

#### Epigenetic Annotation of Non-Human Genomes Using GENA-LMs

While the results discussed previously demonstrate the high efficacy of fine-tuned GENA-LM models in reconstructing and analyzing genomic features such as promoters or protein binding sites, task specific training of these models necessitates the availability of experimentally measured data, which is often lacking for non-model species. We hypothesize that features relatively conserved across different species and cell types could be effectively predicted by a model that has been fine-tuned using human (or other reference) data. To explore this hypothesis, we collected experimentally measured profiles of insulatory protein CTCF binding sites, H3K27 acetylation (H3K27ac) histone marks, and promoter activity across various animal species.

We initiated our study by evaluating a human-based promoter activity prediction model on data from seven species: macaque, mouse, rat, dog, zebrafish, chicken, and *C. elegans*. For mammals (macaque, mouse, rat, and dog), the model’s performance closely mirrored that observed with human promoters, achieving an f1 score of approximately 0.95, despite the absence of species-specific data during the model’s fine-tuning (Fig. 3, A). The evaluation score decreased by 10 pt when applying the same model to phylogenetically more distant vertebrate species such as chicken and zebrafish, but it still achieved a relatively high F1 score of around 0.85. However, for more distantly related species like the fruit fly or the flatworm *C. elegans*, the model’s performance dropped significantly, resulting in an f1 score of approximately 0.7. These results suggest that closely related mammals share similar promoter grammars, which can be captured by training on a human dataset, while more distant species exhibit gradual modifications in the DNA encoding of their promoters.

**Fig. 3.**
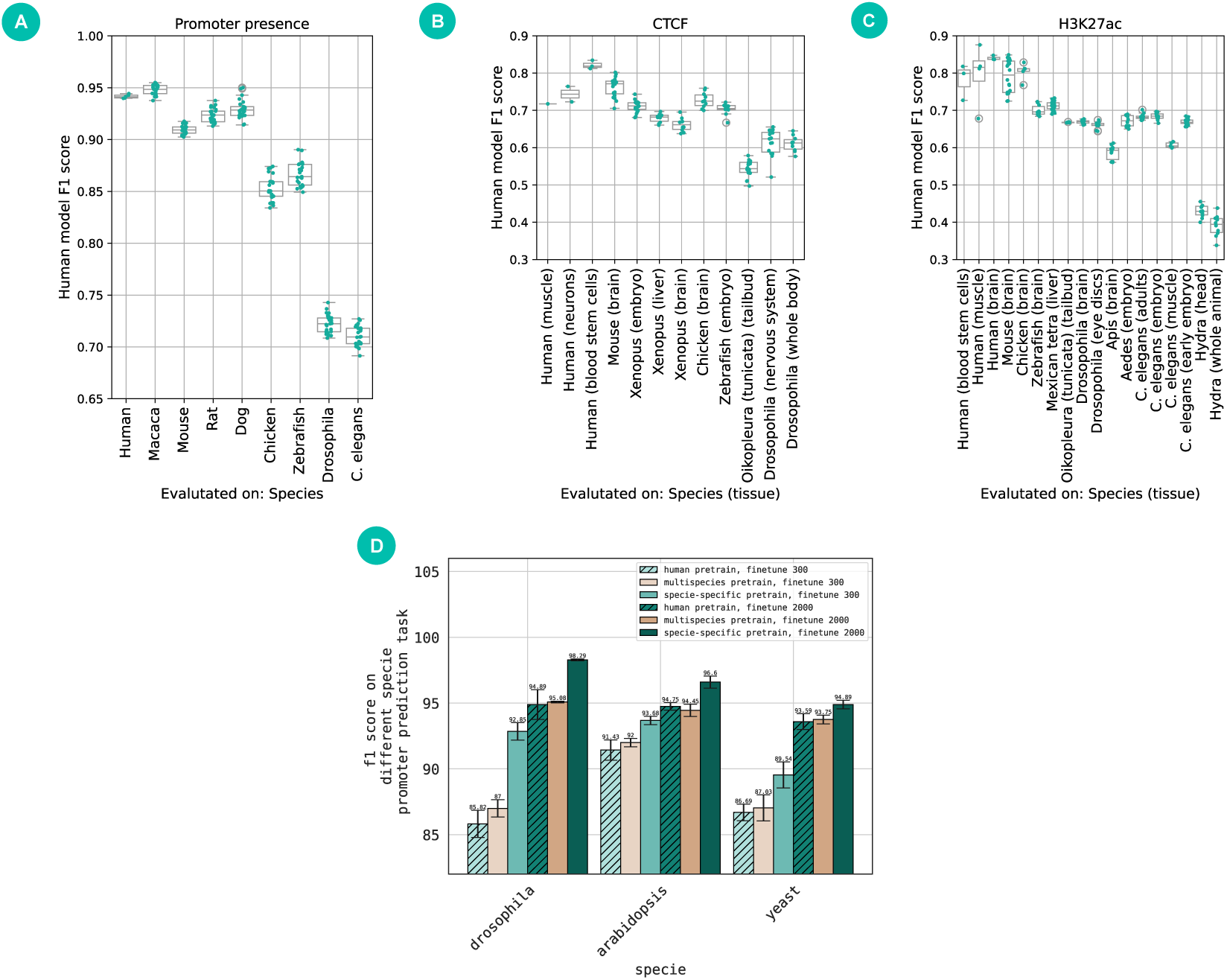
GENA-LMs demonstrate generalisation across species. **A-C**. GENA-LM fine-tuned on human promoters (A), CTCF (B) or H3K27 (C) binding sites evaluated on different species. **D**. Effect of multi-species vs species-specific pertaining on promoter activity prediction.

Then *gena-lm-base-t2t* was trained to differentiate between genomic sites with H3K27ac and CTCF binding sites and those without binding on human data, achieving f1 scores between 0.7 and 0.8 on held-out human sequences (Fig. 3, B-C). Applying this model to other species revealed that its performance correlates with the evolutionary distance between the target species and humans. For CTCF binding prediction, the model demonstrated similar performance in closely related species, such as mice, with a gradual decrease in other vertebrates and a more significant drop in invertebrate species such as *Drosophila* (Fig. 3, B-C).

While similar cross-species analyses for CTCF can be performed using a known motif, this is infeasible for H3K27ac due to the lack of any recognized motif. When evaluating the human H3K27ac model across species, we observed a substantial performance decline outside tetrapod species, and for evolutionarily distant species like Hydra f1 score drops to °0.4. The decline in model performance across species may serve as a measure of the interspecies differences in H3K27ac encoding. Thus, the use of GENA-LMs enables explorations into the evolution of sequence grammar. Furthermore, these results suggest that GENA-LMs can be used to infer epigenetic marks from genomic sequences in the absence of experimental data. Although the performance of such inferences is inferior to that of species-and cell type-specific models and may be limited to taxonomically close groups, it presents extensive opportunities for annotating available genomes, such as using the human model to annotate H3K27ac across several thousand available mammalian genomes.

#### Species-specific pre-training improves model quality

Based on the experiments presented above, it is clear that promoter grammar varies significantly across evolutionarily distant species such as humans, flies, yeasts, and plants [40]. As a result, learning species-specific information during pre-training could prove beneficial for accurate promoter presence prediction and other species-specific genomic tasks. However, pretraining species specific GENA-LMs requires sufficient amount of data and computational resources. Thus, it is important to know whether transfer learning, when a model pre-trained on one data distribution is reused for tasks in similar domain, can substantially improve performance. Another viable strategy in this case involves generation of a universal foundational model trained on the mixture of species.

To study generalisation capabilities of GENA-LM models we utilized EPD promoter annotations for yeast, fly, and Arabidopsis species to fine-tune universal, multispecies or species-specific models. We discovered that for all these species, *gena-lm-bert-base-lastn-t2t* pre-trained on human data can be effectively fine-tuned to provide reasonably accurate promoter classification (Fig. 3, D). When comparing datasets with varying promoter lengths (300 bp versus 2 kb), we observed a significant impact of contextual information on prediction accuracy, mirroring the dependency noted in human data. Compared to the model pre-trined on human data, multispecies pre-training proved to be benefitial: in five out of six experiments, we observed performance improvements ranging from 0.2 to 1.5 points when fine-tuning *gena-lm-bert-base-t2t-multi* model. It’s important to note that the *gena-lm-bert-base-t2t-multi* training dataset includes dozens of genomes; hence, the amount of data from each species available during the pre-training phase was limited. Consequently, we pre-trained new taxon-specific models for yeast, flies, and Arabidopsis. Fine-tuning these models to predict promoter activity in their respective taxa resulted in performance improvements of 2-7 points compared to the model pre-trained on human data (Fig. 3, D). Therefore, we conclude that taxon-specific models can significantly enhance DNA annotations by learning the unique DNA grammar of each species during pre-training. We have publicly released these taxon-specific models to facilitate further applications within the selected species.

#### Species classification based on embeddings from GENA-LMs

The universality of GENA-LMs in addressing various biological challenges suggests that the DNA embeddings of the pre-trained model encapsulate significant biological insights. Evolutionary distant species are known to exhibit divergence in their regulatory code and codon usage patterns. If GENA-LMs effectively capture these inherent biological characteristics during pretraining, one would expect that their embeddings could differentiate DNA sequences sourced from varying species without any additional fine-tuning. To evaluate this premise, we curated a set of 27 species spanning diverse taxonomic classifications, ranging from bacteria to humans (refer to Supplementary Table 3). These species also represent a broad spectrum of evolutionary divergence times, spanning from millions to over a billion years (as depicted in Fig. 4, A, B). We then examined the embeddings generated by inputting genomic DNA subsequences into pre-trained GENA-LMs. Our investigations encompassed a range of sequence lengths, beginning with the typical length of shotgun sequencing reads (100 bp) and culminating at 30 kb, a size consistent with reads from third-generation sequencing platforms.

**Fig. 4.**
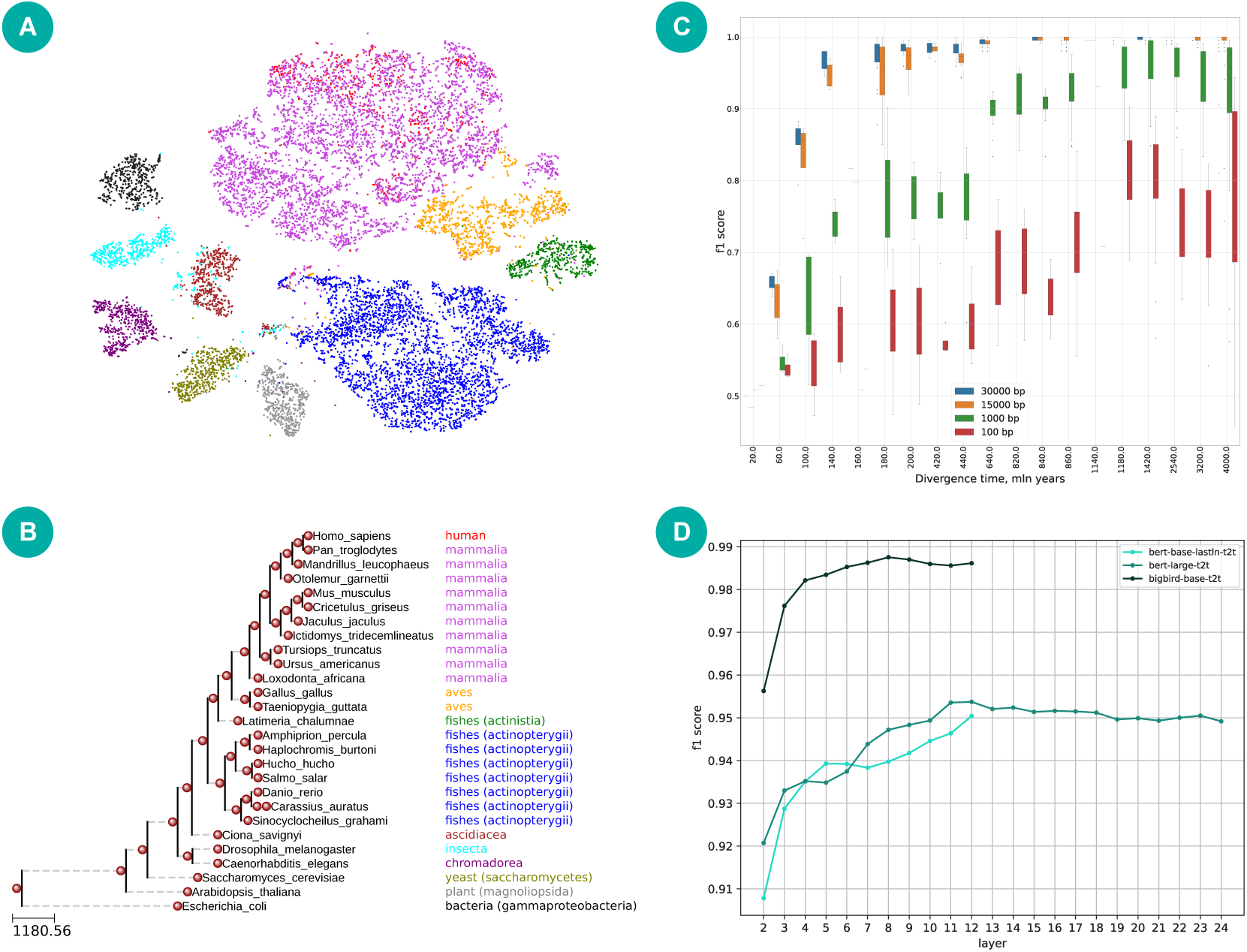
Sequence embeddings from pre-trained GENA-LMs facilitate species classification. **A** and tSNE projections (A) of sequences sampled from 27 species (B), representing a spectrum across the tree of life. **C**. Classification performance for different sequence lengths plotted against divergence time. **D**. Classification performance of embeddings taken from different layers of three models. Data are presented for sequence lengths of 5 kbp (for *gena-lm-bert-base-lastln-t2t* and *gena-lm-bert-large-t2t* ) and 30 kbp (for *gena-lm-bigbird-base-t2t* ).

Initially, we employed tSNE to project sequence embeddings derived from all genomes into a 2D space. This visualization reveals discernible clusters that mirror the phylogenetic relations between the species. Notably, distinct groupings emerged for bacteria, plants, and yeasts, each isolated from the clusters representing animal genomes. Within the realm of animals, we could discriminate invertebrate species and various vertebrate classes. This evidence underscores that GENA-LM embeddings encapsulate nuances allowing for the differentiation of species based on their genomic sequences.

To deeper study these capabilities, we employed a Gradient Boosting algorithm for each of the 27 species pairs. Our aim was to achieve binary species classification leveraging the sequence embeddings.

The data in Fig. 4, C show the richness of information contained in GENA embeddings, enabling species differentiation based on their genomic DNA subsequences. The accuracy of classification is influenced by both the divergence time and the length of the input sequence, with the latter exerting a more pronounced effect. For species that are closely related (with divergence times ≤ 20 MYA), classification accuracy remains constrained (f1 score ≈ 0.7). However, for species diverging around 60–100 MYA-equivalent to the evolutionary separation among all mammalian species-employing the model that accepts longer sequence inputs boosts our classification capability, yielding an f1 score exceeding 0.8. Remarkably, for extensive divergence times (≥ 200 MYA, reflecting the era of the last common ancestor of vertebrates), the classification’s precision approaches perfection.

We next evaluated the classification efficacy of sequence embeddings derived from various layers and architectures of GENA-LMs. Across all models, embeddings sourced from the initial layers consistently delivered subpar performance. This performance incrementally improved, peaking around layers 9 to 12. Notably, for both *gena-lm-bert-large-t2t* (comprising 24 layers) and *gena-lm-bigbird-base-t2t* (with 12 layers), a minor performance dip was observed when utilizing embeddings from the final layers. This trend resonates with prior studies in Natural Language Processing (NLP) [41] and protein modeling [42]. Such studies have posited that in transformer-based language models, the terminal layer embeddings encapsulate information tailored to the specific model training task. In contrast, embeddings from intermediary layers are more versatile, proving advantageous in knowledge transfer for tasks not explicitly addressed during the pre-raining phase. The depth of the layer is also indicative of the abstraction degree of the representations. While preliminary layers prioritize local level representations, the advanced layers capture intricate global features, such as binding sites and contact maps [42].

Collectively, our findings demonstrate that the sequence embeddings from pre-trained GENA-LMs encapsulate abundant biological insights, enabling the resolution of genomic challenges without the necessity for fine-tuning.

### 2.5 GENA-LM-based web-service

Given the demonstrated potential of GENA-LMs in various genomic tasks, we aim to extend their accessibility through GENALM-Web, a web-service designed for sequence annotation using DNA language models (Fig. 1, C). GENALM-Web incorporates several downstream tasks developed in this paper, such as promoter activity prediction, chromatin annotation, splice site inference, and enhancer activity prediction for *Drosophila* sequences. Key features of the web-service include the capability to handle exceptionally long inputs (up to 1 Mb), utilize extensive contextual information, and conduct token importance analysis in real-time. This last feature allows users to identify sequence regions responsible for specific features, even if these regions are located at distant genomic locations. The web-service is accessible at https://www.dnalm.airi.net, and documentation is available at service pages and in the supporting manuscript [43].

### 2.6 Handling even longer sequences with recurrent memory

While the integration of sparse attention techniques and BPE tokenization in GENA-LMs has substantially expanded the permissible DNA input length, the current limit (about 36 kb) may not sufficiently capture certain biological dependencies. Notably, the prediction of chromatin interactions [7], enhancer-promoter associations [4], gene expression [8], and other genomic phenomena necessitate the processing of contexts that extend beyond 30 kb. Additionally, our empirical analyses show improvements in promoter and splice site predictions as the context size expands from 512 to 4096 tokens (Section 2.2). This indicates the potential benefits of further enhancing sequence length for these biological tasks.

To enhance the input capacity of GENA-LMs, we incorporated recurrent memory mechanisms. The Recurrent Memory Transformer (RMT) has been demonstrated as an efficient, plug-and-play method to handle extended input sequences using pre-trained Transformer models [30]. In this recurrent strategy, the input sequence is partitioned into segments which are processed one after the other (Fig. 5, A). Special memory tokens are introduced to each segment to pass information between consecutive segments, allowing them to use information from all previous segments. Thus, the entire pre-trained Transformer effectively functions as a single recurrent unit.

**Fig. 5.**
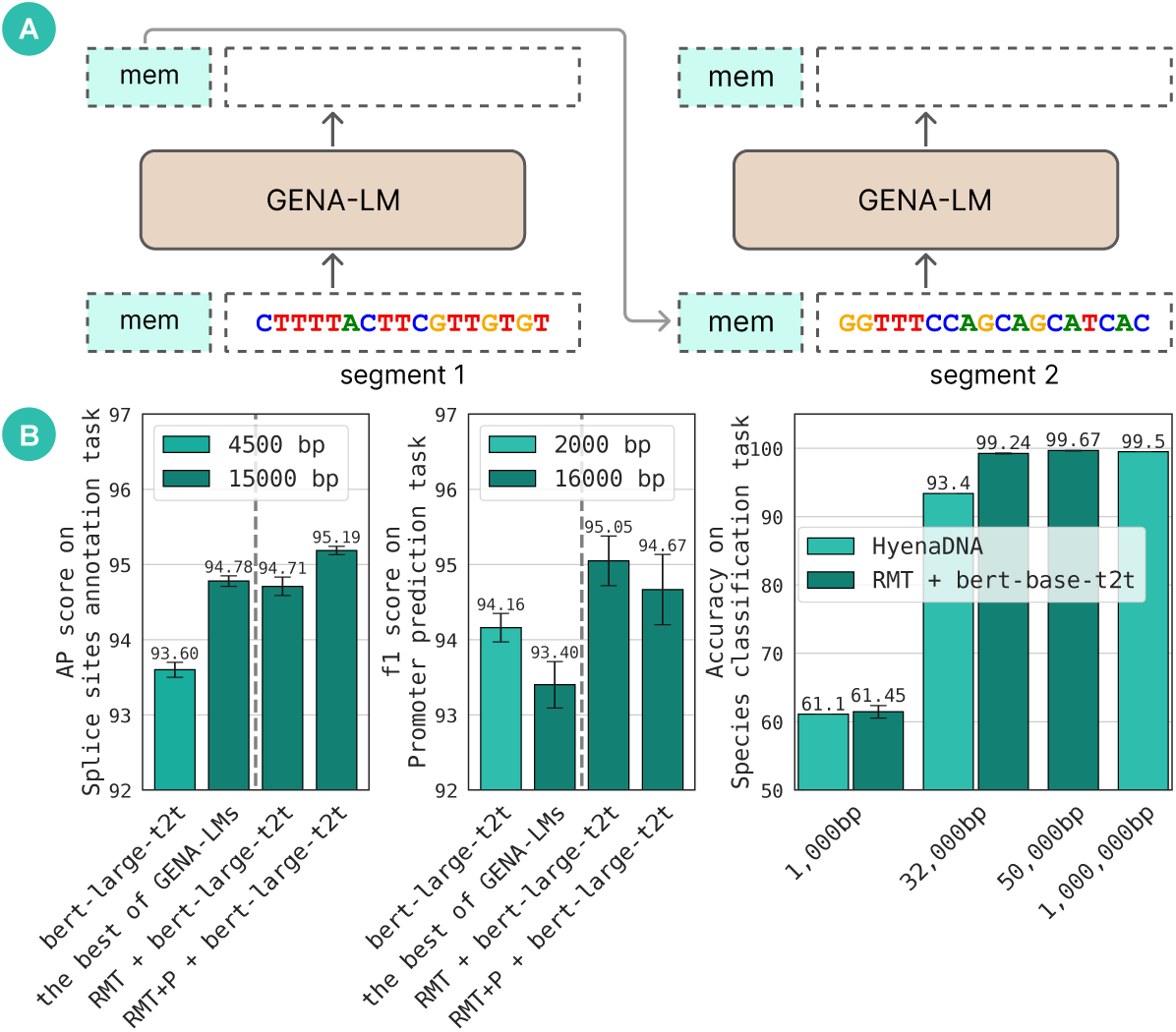
Leveraging Recurrent Memory to enhance the input capacity of GENA-LM models yields improved performance in downstream tasks. **A**. The Recurrent Memory Transformer (RMT) architecture. A vocabulary of the model is augmented with a memory token denoted as *mem* on the diagram. Memory augmented model is fine-tuned to write relevant information in memory tokens and pass it to subsequent segments. **B**. The augmentation of GENA-LM with RMT with 3x (left), 8x (center), and 50x (right) larger sequence lengths. Models with memory achieve superior results in splice site annotation and promoter prediction tasks when compared to all other GENA-LMs, including those utilizing sparse attention (Wilcoxon test P-value *≤* 0.043 in all comparisons). AP - average precision. On the species classification task, RMT with GENA-LM outperforms the HyenaDNA model designed for long sequences. RMT+P refers to models that have not only been fine-tuned with RMT, but also pre-trained with it.

RMT can be optionally incorporated during pre-training, enabling the model to learn the use of memory tokens at this stage. Alternatively, memory tokens can be introduced during the fine-tuning phase, using a model that was pre-trained without RMT. To assess these two training strategies and determine the optimal approach, we conducted pre-training experiments with *gena-lm-rmt-base-t2t* and *gena-lm-rmt-large-t2t* models. During pre-training, we evaluated how RMT augmentation influences masked language modeling accuracy. As illustrated in Supplementary Fig. 8, RMT augmentation enhances accuracy for the base-size model when provided with 8–10 segments of contextual information. However, the large-size model pre-trained without RMT augmentation achieves a substantially better score. Extending the large-size model’s input with RMT does not improve the score (Supplementary Fig. 8). These findings corroborate our previous observation that *gena-lm-bert-large-t2t* outperforms *gena-lm-bigbird-base-sparse* in the MLM task Fig. 1, F, despite the latter’s longer input length. Therefore, we propose that long-range dependencies beyond 4 kb are not necessary for accurate masked token prediction. However, these dependencies are important for downstream tasks.

For a comparative evaluation between RMT and other GENA models, we focused on tasks with inputs of moderate length (15–16 kb), which can be processed by sparse models. The *gena-lm-bert-large-t2t* model, when integrated with RMT, was fine-tuned on sequences of 16 kb for promoter prediction and 15 kb for splice site prediction. Inputs were divided into segments, with each segment comprising approximately 512 tokens or about 4.5 kb. These segments also included memory tokens as part of the input, with 10 memory tokens used for each task. When contrasted with the original *gena-lm-bert-large-t2t* model, the sequence length processed by the *gena-lm-bert-large-t2t* + RMT increased substantially: from 3 to 8 times (rising from 4.5 kb to 15 kb for the splice site prediction and from 2 kb to 16 kb for other tasks).

The expansion in input length significantly enhanced the performance of the *gena-lm-bert-large-t2t* model, as depicted in Fig. 5, B. Notably, models employing the Recurrent Memory Transformer (RMT) outperformed all other GENA-LMs, including those sparse variants of GENA-LM. While these sparse models can accommodate the input lengths featured in the aforementioned tasks, they have fewer parameters compared to *gena-lm-bert-large-t2t*. Thus, RMT allows combining models with the higher number of parameters and longer sequence inputs, achieving the best performance on the common biological tasks. Furthermore, RMT has no limit in a sequence length and could be used for even longer sequences. Sparse GENA-LMs, on the other hand, are limited to the lengths on which they were trained.

We next used splice sites and promoter activity prediction tasks to benchmark the effects of the RMT application during pre-training. This benchmark yields mixed results: for promoter activity prediction, limiting the RMT approach to the fine-tuning stage does not diminish performance compared to its use at both the pre-training and fine-tuning stages. However, for splice site annotation, pre-training with RMT proved beneficial, improving performance by approximately 0.5 points.

Recently, the HyenaDNA team introduced a specific benchmark for DNA language models that process long input sequences [34]. The authors demonstrated that HyenaDNA can classify five mammalian species (human, mouse, lemur, pig, and hippo) based on their genomic sequences. Compared to the classification of a broader range of species using GENA-LM embeddings, as previously described, the species in this benchmark are phylogenetically closer, making classification more challenging. Original research [34] illustrated that classification accuracy is heavily dependent on the input DNA length, which increases gradually from 61.1 to 99.84 as sequence length scales from 1 kb to 1 Mb.

We compared HyenaDNA results with *gena-lm-bert-base-t2t* augmented with RMT. For sequence inputs of 1 kb, the classification accuracy of both models was relatively low (61.45 ± 0.91 for GENA and 61.1 for HyenaDNA). A significant enhancement in classification accuracy was observed when the sequence length was increased to 32 kb, with *rmt+gena-lm-bert-base-t2t* achieving 99.24 ± 0.06, thereby surpassing the performance of HyenaDNA (93.4), as shown in Fig. 5, right. Further extending the sequence length to 50 kb elevated the classification accuracy of *rmt+gena-lm-bert-base-t2t* to 99.67 ± 0.059, exceeding the accuracy HyenaDNA attained with 1000 kb sequences. This indicates that, within this experimental framework, RMT and the associated model architecture extract and leverage information from extended DNA sequences more efficiently than other technologies designed for processing long input sequences such as Hyena layers underlying HyenaDNA.

## 3 Discussion

Transformer architectures have garnered significant interest across diverse research domains, including genomics. They consistently achieve exemplary results in various biological tasks such as deciphering gene expression regulation in mammals [8] and *E. coli* [44], predicting phenotypes from gene expression [45, 46], deducing DNA methylation [47], and filling in missing genotypes [48], to name a few. Nevertheless, the challenge lies in training task-specific models for each distinct biological question. This process demands substantial time and resources. DNABERT, DNABERT-2, and similar foundational DNA models such as BigBird and Nucleotide Transformer provide a solution by offering a platform for refining universally applicable models without starting from scratch. The Nucleotide Transformer v2 [31] has incorporated Rotary Embeddings and Gated Linear Units paired with swish activations, distinguishing it from its predecessor. This model has an input size of 12 kb. DNABERT v2[17], while drawing upon the foundational DNABERT architecture, expands in terms of model parameters and employs BPE tokenization. However, its sequence input length remains below 4000 bp. Contrarily, HyenaDNA [34] introduces a novel architecture capable of handling vast DNA sequences, extending up to 1 million base pairs. Yet, benchmark results suggest an inverse relationship between HyenaDNA’s performance and the input size used during its training, as noted by [31]. Moreover, our benchmarking GENA-LMs augmented with RMT against HyenaDNA in species classification task indicates better performance of the former model.

A unique feature of HyenaDNA is its decoder-only configuration. Unlike the encoder-centric GENA-LMs, HyenaDNA doesn’t generate sequence embeddings directly. Instead, it produces DNA sequences, making the derivation of class labels (for classification purposes) or quantitative targets (for regression) from its outputs a complex task. To predict specific DNA states with the HyenaDNA model, the authors utilized a DNA-alphabet encoding, obliging the model to understand this biologically unrelated nucleotide sequence interpretation.

We introduce GENA-LMs, a collection of open-source models boasting the most extensive input capacity among all accessible DNA transformers^4^. The GENA-LM collection encompasses a spectrum of publicly accessible architectures, catering to researchers by offering tailored solutions for unique challenges. Moreover, GENA-LMs include several taxon-specific models that can improve performance in species-specific setups. Our rigorous benchmarking affirms that GENA-LMs not only surpass earlier pre-trained models but occasionally even rival the precision of task-specific convolutional neural networks.

In our comparison of various GENA-LMs, we investigated the influence of context length and the total number of model parameters on predictive accuracy. We found that the optimal balance between these two factors varies depending on the specific task. For instance, an extended context is vital for predicting promoter activity or deciphering widespread histone mark distributions, as previously indicated by [5]. However, for certain tasks, a more concise context is adequate, making it more advantageous to augment the model’s parameter count. The broad spectrum of GENA-LMs available offers researchers the flexibility to select a model best suited for their particular objective.

While GENA-LMs accommodate extensive input sizes, they occasionally fall short of the lengths required for peak accuracy in specific biological tasks. For example, research has shown that gene expression can be influenced by variants situated hundreds or even millions of base pairs distant from the promoter. This can be attributed to processes such as loop extrusion [49] and other 3D-genomic mechanisms [50]. There are several strategies to address this constraint in GENA.

First, the RMT technique facilitates the processing of extensive sequence inputs using powerful models with a large number of parameters. Our benchmarks reveal that this approach delivers superior results for tasks where the biological signal spans a lengthy context. Notably, unlike transformer layers that exhibit a quadratic memory dependence on the number of tokens, the computational resources needed for RMT training and inference scale linearly with sequence length. RMT can be integrated with GENA-LMs not only during the fine-tuning phase of downstream tasks but also throughout the MLM pre-training stage. This could enhance learning operations on extended sequences, particularly for downstream task datasets that are of limited size. Furthermore, RMT models pre-trained on multiple segments can be utilized for a greater number of segments during inference [30]. As such, RMT models are versatile enough to address a variety of downstream tasks, even for teams without access to cutting-edge computational infrastructure.

Second, the 3D proximity of chromatin can be determined using specialized models [7]. This information can then be directly incorporated into transformer models, enabling them to capture long-range associations between functional genomic elements.

One limitation of GENA-LMs arises from the granularity imposed by the use of BPE tokenization, which confines predictions to specific tokens. To overcome this, exploring alternative DNA tokenization methods and developing low-level nucleotide embeddings could offer solutions for certain applications.

Beyond merely predicting specific biological signals, we demonstrate that GENA-LMs can also be harnessed to decipher and understand the functions of sequences underpinning these signals. An analysis of token importance revealed that GENA-LM accurately detected motifs corresponding to known transcription factors. Furthermore, it pinpointed transcription factor binding sites crucial for specific histone modifications. There exists an array of factors that modify histones, termed histone ”writers”, many of which are cell-type specific. Determining these factors and their corresponding genomic binding sites is a complex endeavor. In this context, we illustrate how GENA-LMs can aid in this task by discerning motifs deemed ”essential” for a specific histone mark within a particular cell type. However, it’s pivotal to approach this method judiciously. The presence of enriched motifs may only indicate an association rather than a direct causal relationship. As an instance, while the enrichment of recognized activator factor motifs within H3K4me3-important tokens aligns with the understood biological roles of these factors, the presence of motifs specific to neural or dermal transcription factors within H3K27me3-important tokens in lymphoid K562 cells likely doesn’t signify a direct causal role of these proteins in establishing the repressive H3K27me3 mark. We posit that these factors’ targets were suppressed in blood lineage progenitors, implying that the enrichment of their motifs is a reflection of the developmental trajectory of these cells.

To sum up, our study provides compelling evidence that large language models trained on DNA sequences have the capability to generate useful biological insights. This not only presents an innovative method for solving an array of genomic challenges but also forges a pathway for a more nuanced understanding of genetic data. The transformative impact of language models has already been witnessed in protein biology, where they have brought about remarkable progress in predicting protein properties and engineering novel peptides with tailored functions ([51–53]). This is indicative of the potential these models hold, suggesting that their capabilities go beyond mere sequence analysis. With the exponential increase in multi-omics data—spanning genomics, transcriptomics, proteomics, and metabolomics—it’s imperative to have advanced analytical tools that can seamlessly integrate and interpret these vast and complex datasets. Language models, as demonstrated by our findings, appear poised to fill this role. As the nexus between computational techniques and biology strengthens, it is foreseeable that language models will be pivotal in ushering in a new era of DNA-based technologies.

## 4 Methods

### 4.1 Datasets

#### 4.1.1 Genomic datasets for language model pre-training

##### Dataset sources

Human T2T v2 genome assembly was downloaded from NCBI (acc. GCF 009914755.1). Genomic datasets used to train multispecies models were downloaded from ENSEMBL release 106^5^. The list of species is provided in Supplementary Table 4. For the 1000-genome dataset, we used gnomAD v. 3.1.2 data. For taxon-specific models, we used the following resources:

1. Arabidopsis models: Data were obtained from [54] and contains chromosome-level genomes of 32 *A. thaliana* ecotypes.
2. Yeasts model: Data were obtained from [55] and includes telomere-to-telomere assemblies of 142 yeast strains.
3. *Drosophila* model: Data were obtained from Progressive Cactus alignment of 298 drosophilid species generated by [56].

##### Genomic datasets preprocessing

To prepare genomic datasets for our training corpus, we processed each record in the genomic fasta files. We excluded contigs with the substring ”*mitochondrion*” in their identifiers and those shorter than 10 kb. From the remaining sequences, we divided them into ’sentences’ spanning 500 to 1,000 base pairs (bp) — the sentence length being randomized — and compiled ’documents’ with 50 to 100 consecutive sentences. This approach follows the data processing in BigBird [21]. Data augmentation incorporated reverse-complement sequences, and we applied a stochastic shift for some documents to include overlapping genomic sequences.

For the 1000-genome SNP augmentation, nucleotide substitution was executed, replacing reference alleles with alternative ones sourced from individual samples of 1000-genome cohort. In order to maintain the haplotype structure, each sample was processed individually. This meant that for every genomic region, multiple sequences were derived, each resulting from swapping reference alleles with sample-specific alternative variants from singe individual. We limited our focus to genomic regions where the proportion of positions with a noted variant for a given sample exceeded 0.01. No allele frequency filter was applied.

##### Train and test split

For our initial models, *bert-base* and *bigbird-base-sparse*, we hold out human chromosomes 22 (CP068256.2) and Y (CP086569.2) as the test datasets for the masked language modeling task. In contrast, for subsequent models, identifiable by the ”t2t” suffix in their names, we hold out human chromosomes 7 (CP068271.2) and 10 (CP068268.2) for testing. All remaining data was used for training.

Models focusing exclusively on human data were trained using the pre-processed Human T2T v2 genome assembly combined with its 1000-genome SNP augmentations, totaling approximately ≈ 480 × 10^9^ base pairs. On the other hand, multispecies models incorporated both the human-only and multispecies data, aggregating to roughly ≈ 1, 072 × 10^9^ base pairs.

The data splitting strategy for downstream tasks was anchored to methodologies previously described in literature relevant for each particular downstream task. Comprehensive specifics for each task are provided in their respective dedicated sections.

##### Sequence tokenization

We employed Byte-Pair Encoding (BPE) tokenization [57] for our models, setting the dictionary size to 32,000 and initializing with a character-level vocabulary comprised of [’A’, ’T’, ’G’, ’C’, ’N’]. Our study utilized two distinct tokenizers:

1. The first, trained exclusively on the human T2T v2 genome assembly, is denoted as ’T2T split v1’ in Table 4.
2. The second tokenizer, trained on a mixture of human-only and multispecies data sampled equally, is labeled ’T2T+1000G SNPs+Multispecies’.

Both tokenizers incorporate special tokens: CLS, SEP, PAD, UNK, and MASK. Notably, the ’T2T+1000G SNPs+Multispecies’ tokenizer integrates a preprocessing step to manage extensive gaps: sequences with over 10 consecutive ’N’ characters are consolidated into a singular ’–’ token.

#### 4.1.2 Downstream task datasets

A concise overview of the dataset parameters for downstream tasks is presented in Table 3. A comprehensive description follows.

**Table 3.** Parameters of downstream tasks datasets.

| Downstream task | Input length, bp | Number of targets | Task |
| --- | --- | --- | --- |
| Promoters prediction (300) | 300 | 2 | classification |
| Promoters prediction (2,000) | 2,000 | 2 | classification |
| Promoters prediction (16,000) | 16,000 | 2 | classification |
| Splice site prediction | 15,000 | 3 per token / bp | multi-class classification |
| Drosophila enhancers prediction | 249 | 2 | regression |
| Chromatin profiling (1,000) | 1,000 | 919 | multi-label classification |
| Chromatin profiling (8,000) | 8,000 | 919 | multi-label classification |
| Polyadenylation sites prediction | 443 | 1 | regression |

##### Promoters prediction

For the task of predicting promoters, we sourced human sequences located upstream of TSS (transcriptional start sites) from the EPDnew database^6^. Sequences of lengths 300 bp, 2,000 bp, and 16,000 bp were extracted, with each dataset being processed and assessed independently. For negative samples generation, we randomly selected genomic locations outside promoter sequences, ensuring that negative and positive samples do not overlap for maximum promoter length (16 kb). The entire dataset was segregated by sequence into training, validation, and testing sets. The objective of this task is a binary classification: determining the presence or absence of a promoter within a given region.

##### Splice site prediction

To predict splice donor and acceptor sites, we replicated the dataset from [58], utilizing the original scripts provided by the authors. We adhered to the same training and testing splits as outlined in [58]. In this dataset, a central 5,000 bp target region is bracketed by 10,000 bp of context, with 5,000 bp on each side. Splice site annotations within the target region are aligned to token positions. Tokens overlapping with either splice-donor or splice-acceptor sites are designated as positive samples for their respective splicing annotation class. Subsequently, both the target and its context were tokenized independently. If the combined length diverged from the model’s input size, adjustments were made through either padding or truncation. In the event of truncation, sequences furthest from the target region’s midpoint were first removed. We demarcated the context and target sequences using SEP tokens. Through this procedure, the target’s size matched the model’s input token count. However, the computational loss did not account for tokens representing either context or padding. This challenge is a multi-class, token-level classification task encompassing three categories: splice donor, splice acceptor, and none.

##### Drosophila enhancers prediction

Candidate sequences, along with their associated housekeeping and tissue-specific activity in *Drosophila* cells, were sourced from the Stark Lab repository^7^. These datasets are partitioned into training, validation, and testing sets, consistent with those used for training the Deep-STARR model [32]. The task at hand involves a two-class regression, wherein each 249-bp sequence is predicted to produce two continuous scores: one for housekeeping enhancer activity and another for developmental enhancer activity.

##### Chromatin profiling

We gained the DeepSEA dataset [33] from its original repository^8^. This dataset outlines the chromatin occupancy profiles of various genomic features, encompassing histone marks, transcription factors, and DNAse I hypersensitivity regions. The dataset comprises DNA sequences of 1,000 bp, with a central 200 bp target region flanked by 400 bp contexts on either side. Each feature’s occupancy is quantified over this 200 bp target. Additionally, we trialed an expanded context of 7,800 base pairs (yielding a total input length of 8,000 bp). To elongate the DNA context, we aligned the input DNA segments to the hg19 genome using *bwa fastmap*. Surrounding sequences at mapped sites were then harvested. Sequences that either failed remapping or aligned too proximate to a chromosome’s terminus to permit extension were omitted, though these comprised less than 1% of the dataset. Our partitioning for training, validation, and testing adhered to the divisions presented in the original DeepSEA dataset. The challenge is a multi-label classification, with class count reflecting the unique epigenetic profiles identified in DeepSEA (919 in total).

##### Polyadenylation sites prediction

For predicting polyadenylation sites, we employed the APARENT dataset [59] (available at^9^). This dataset characterizes the frequency with which transcription machinery recognizes specific nucleotide sequences as polyadenylation signals. Utilizing the scripts published by the authors, we extracted the target values and delineated the training and testing datasets. Furthermore, we retrieved APARENT predictions (noted under the field *iso pred* ) to gauge the performance of the APARENT model. We tokenized the sequences from both upstream and downstream segments of the 5’-untranslated regions individually, and they were demarcated using a SEP token. This study focuses on regression analysis targeting 256 bp sequences.

##### Nucleotide Transformer Dataset

The dataset and literature scores were obtained from the Nucleotide Transformer (NT version 2 scores) [31].

##### HyenaDNA species classification dataset

The dataset was reconstructed based on the description provided in [34]. Genomes from 5 species (human, lemur, mouse, pig, hippo) were downloaded from NCBI (Ref-Seq assemblies GCF 000001405.40, GCF 020740605.2, GCF 000001635.27, GCF 000003025.6, GCF 030028045.1 respectively). Four chromosomes (chromosomes 1, 3, 12, and 13) were used for models evaluation, other chromosomes were utilized during training. We sampled sequences from chromosomes randomly, using the uniform distribution. We used a 5-way classification and reported top-1 accuracy. For each task length, we collected a total of 50 000 DNA subsequences from each species, ensuring a comprehensive dataset for our analysis.

### 4.2 Models architecture and training

#### 4.2.1 DNA language models based on transformer architecture

We trained and expanded upon several transformer models, drawing inspiration from both BERT [14] and BigBird [21] architectures. These adapted models are consistently referred to as GENA-LM throughout this manuscript. Key distinctions between these architectures can be found in Table 4. A comprehensive breakdown of parameters and specific combinations for each model is available in Supplementary Table 5. Additionally, we enhanced BERT-based models with Pre-Layer normalization [60]. In instances where the layer normalization is applied even to the final layer output, it is distinctly mentioned as *lastln* in the model names. For precise parameter details, refer to Supplementary Table 5.

**Table 4.** Overview of the GENA-LM Foundational DNA Language Models. This table delineates the specifications of pre-trained GENA-LM models, highlighting variations in pre-training data, layer count, attention type, and sequence length. Models archived on the HuggingFace model hub adhere to a consistent naming convention, prefixed by AIRI-Institute/gena-lm-. Models based on the BERT architecture utilize Pre-Layer Normalization [60], with lastln indicating the application of layer normalization to the output of the terminal layer. ’T2T split v1’ alludes to initial experiments using a non-augmented T2T human genome assembly split. The term ’1KG’ is shorthand for 1000G SNPs augmentations, while ’M’ denotes the inclusion of Multispecies data. The designations ’DS Sparse’ and ’HF Sparse’ are references to the DeepSpeed sparse attention and HuggingFace BigBird implementations, respectively. The abbreviation ’RoPE’ signifies the adoption of rotary position embeddings [62] as an alternative to BERT’s absolute positional embeddings. The models were structured with either 12 (denoted BERT-12L) or 24 (denoted BERT-24L) layers, comprising 110M and 336M parameters, respectively.

| Model | Architecture | Maximum seq len, tokens (≈ bp) | Tokenizer data | Training data |
| --- | --- | --- | --- | --- |
| DNABERT | BERT-12L | 512 (512) | 3,4,5,6-mer | GRCh38.p13 |
| GENA-LM models: |  |  |  |  |
| *bert-base* | BERT-12L | 512 (4,500) | T2T split v1 | T2T split v1 |
| *bert-base-t2t* | BERT-12L | 512 (4,500) | T2T+1KG+M | T2T+1KG |
| *bert-base-lastln-t2t* | BERT-12L | 512 (4,500) | T2T+1KG+M | T2T+1KG |
| *bert-base-t2t-multi* | BERT-12L | 512 (4,500) | T2T+1KG+M | T2T+1KG+M |
| *bert-base-t2t-yeast* | BERT-12L | 512 (4,500) | T2T+1KG+M | Yeast |
| *bert-base-t2t-fly* | BERT-12L | 512 (4,500) | T2T+1KG+M | *Drosophila* |
| *bert-base-t2t-athaliana* | BERT-12L | 512 (4,500) | T2T+1KG+M | *A. thaliana* |
| *bert-large-t2t* | BERT-24L | 512 (4,500) | T2T+1KG+M | T2T+1KG |
| *bigbird-base-sparse* | BERT-12L, RoPE<br>DS Sparse Att | 4,096 (36,000) | T2T split v1 | T2T split v1 |
| *bigbird-base-sparse-t2t* | BERT-12L, RoPE<br>DS Sparse Att | 4,096 (36,000) | T2T+1KG+M | T2T+1KG |
| *bigbird-base-t2t* | BERT-12L<br>HF Sparse Attention | 4,096 (36,000) | T2T+1KG+M | T2T+1KG |

All models were pre-trained using the masked language modeling (MLM) objective. During this process, the sequence was tokenized and flanked by the special tokens, CLS and SEP. In alignment with the BERT pre-training methodology, 15% of the tokens were randomly selected for prediction. Among these, 80% were replaced with MASK tokens, 10% were swapped with random tokens, and the remaining 10% were retained unchanged. Training extended for 1-2 million steps, utilizing a batch size of 256 and operated on 8 or 16 NVIDIA A100 GPUs. We employed the FusedAdam implementation of the AdamW optimizer [61], made available through Nvidia Apex^10^. The initial learning rate was set at 1 × 10^−4^, including a warm-up phase. For most models, we adopted a linear learning rate decay, but in cases where pre-training diverged, we manually adjusted the learning rate.

#### 4.2.2 GENA-LM fine-tuning

In our standard procedure, we tokenize input sequences and prepend and append them with the service tokens CLS and SEP, respectively. To ensure compatibility with the model’s input requirements, sequences are either padded or truncated as needed. For datasets necessitating specialized tokenization, the specific preprocessing steps are detailed in the relevant dataset section.

Tokenized sequences were provided as inputs to downstream models. These models utilized one of the pre-trained GENA-LM architectures, augmented with a single fully-connected output layer. The dimensions of this layer are denoted by (hidden size, target size). Here, hidden size refers to the hidden unit size specific to the GENA-LM model (refer to Supplementary Table 5), while target size is specified in the description of each downstream task dataset discussed earlier. For single-label, multi-class classification tasks, we implemented a softmax activation function on the final layer, paired with cross-entropy loss. In contrast, multilabel, multi-class classification tasks employed a sigmoid activation function on the last layer, combined with a binary cross-entropy with logits loss. Regression tasks did not necessitate any activation function on the last layer and utilized mean squared error as the loss function. To address sequence classification and regression tasks, we used the hidden state of the CLS token from the final layer. Meanwhile, for token-level classification tasks, such as splice site prediction, all hidden states from the ultimate layer were employed. Both the weights of the final fully connected layer and the parameters of the entire GENA-LM were fine-tuned during this process. Learning rate warm-up [63] was consistently applied across all tasks. The optimal number of training and warm-up steps was determined empirically for each individual task.

#### 4.2.3 GENA-LM fine-tuning with recurrent memory

The recently introduced Recurrent Memory Transformer (RMT) presents a novel approach to extend the context length of pre-trained models [29]. Unlike traditional transformers, which exhibit quadratic computational complexity in their attention layers, the RMT employs a recurrent mechanism to efficiently manage elongated sequences. This recurrent design ensures constant memory consumption and linear computational scaling with context length. To process input, the RMT divides the sequence into distinct segments, processing them in a sequential manner. Special memory tokens are integrated into the input of each segment. For a given segment, the outputs linked to its memory tokens are subsequently utilized as input vectors for memory tokens for the succeeding segment. By this method, a multi-layer transformer, such as the pre-trained GENA-LM, functions as a single recurrent cell, addressing one segment at a time.

For both promoter and splice site prediction tasks, we segmented the input sequence into units, with each containing 512 tokens (approximately 4.5 kb). The initial 10 tokens of every sequence were allocated for memory tokens. Segments were processed in a sequential manner, where outputs from the memory tokens of one segment are used as the input memory tokens of the subsequent segment. During the training phase, gradients were allowed to propagate from the final segment to the initial one through these memory tokens. We did not impose any restrictions on the number of unrolls in backpropagation through time (BPTT), allowing gradients to flow uninterrupted from the final to the initial segment. The initial states designated for memory tokens were randomly initialized, and further refined during the fine-tuning process. For the task of promoter prediction, we restricted loss computation to only the last segment. Conversely, for splice site prediction, the loss was determined for every individual segment. The training employed the AdamW optimizer and learning rates of {1e-04, 5e-05, 2e-05}. With a batch size set at 128, the training was terminated when there were no discernible improvements in validation scores. The results for the promoter prediction task are presented as averages over five folds. Meanwhile, the splice site prediction task results are averages across three runs, each employing a distinct random initialization. Training scripts are accessible within our provided codebase.

For the species classification task, we used *gena-lm-bert-base-t2t* model that has been augmented with RMT (8 segments) during the pre-training phase. The processes of fine-tuning were enhanced through the application of a curriculum learning strategy. This meant that our initial step included fine-tuning the model on DNA subsequences of 1000 bp in length (single segment). Following this initial phase, we proceeded to extend the fine-tuning process to handle longer DNA subsequences while using the model weights from the 1000 bp fine-tuned model as initial weights, increasing the challenge to a length of 32 kb (8 segments). Continuing with this progressive training methodology, we further advanced our model’s capabilities by eventually fine-tuning it to efficiently process and analyze DNA subsequences extending up to 50 kb in length (12 segments). This gradual fine-tuning approach, in line with the principles of curriculum learning, facilitated the model in sequentially mastering tasks of escalating complexity, thereby enhancing its analytical precision and performance on genetic classification tasks.

### 4.3 Phylogenetic analysis using GENA-LMs without recurrent memory

For our phylogenetic analysis, we randomly sampled 500 subsequences from each genomic sequence, as detailed in Supplementary Table 3. To ensure representative sampling across entire genomes, the probability of selecting a sequence from a specific chromosome was proportionate to the chromosome’s length. In instances not otherwise specified, we utilized the embedding of the CLS token from the final layer. Sequences shorter than 5 kb were processed using the *bert-large-t2t* model, whereas sequences exceeding this length were analyzed with the *big-bird-base-t2t* model to accommodate the extended context. For classifying species, we employed the HistGradientBoostingClassifier from the sklearn library, retaining its default parameters.

### 4.4 Classification of human promoter mutations

We curated records from the ClinVar database as of July 01, 2024, applying several filters to ensure data quality and relevance. Only records with more than one piece of evidence (i.e. filteringby the *single submitter* field) and no conflicting interpretations of significance (as indicated in the *conflicting interpretations* field) were included. We retained only variants with consequences classified as either benign or pathogenic. To focus on regulatory variants, we excluded variants overlapping exons, as defined by Gencode V45. Additionally, we filtered out variants located on sex chromosomes and mitochondrial variants. From the filtered dataset, we specifically selected those single nucleotide substitutions that overlap with promoters within the 2 kb EPDnew promoter dataset described previously.

For each variant, we selected all overlapping promoters and computed the log odds value as follows: 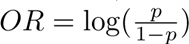, where *p* represents the promoter presence probability derived from the *gena-lm-bert-large-t2t* model, which was fine-tuned on a 2 kb length human promoter dataset. The *OR* was calculated for both the reference sequence and the sequence containing the mutation, and their absolute difference was utilized as the mutation score. If a single mutation overlapped several promoters, the highest score among them was used.

### 4.5 Cross-species epigenetic analysis

For the cross-species analysis of H3K27ac and CTCF binding sites, we collected ChIP-seq data from NCBI, with accession numbers listed in Supplementary Table 7, and uniformly processed them using the MACS3 software. For each dataset, we filtered out peak calls located on scaffolds shorter than 200 kb, and from the remaining data, we randomly selected 2000 binding sites as positive samples. All genomic regions located at least 8 kb away from any positive sample were designated as negative samples, and 2000 negative samples were randomly chosen for each dataset. We next computed the center of each sample and collected 2000 bp of the flanking sequences (± 1000 bp) to provide contextual information.

Each genome was randomly split into five folds, ensuring that sequences from different folds did not overlap and that chromosomes were uniformly distributed across the folds. We fine-tuned the *gena-lm-bert-base-t2t* model using the human SRR10182244 dataset for H3K27ac and the human SRR26329064 dataset for CTCF. When assessing performance on human datasets, we utilized sequences exclusively from one fold (fold 1), which was held out during training. For non-human species, since their data was not included during the human model’s fine-tuning, we performed evaluations on each of the five folds. Consequently, results for each human dataset evaluation are depicted by a single point in Figure 3 A and B, while results for non-human dataset evaluations are represented by five points.

### 4.6 Cross-species promoter inference

We evaluated the *gena-lm-bert-large-t2t* model, which was fine-tuned on human promoter sequences, using data from seven species: macaque, mouse, rat, dog, zebrafish, chicken, and *C. elegans*. For each species, we downloaded promoter sequences from EPDnew and prepared the data into five folds, following the same procedure used for the human dataset. For each species, we conducted 25 evaluation experiments by cross-applying the models trained on each of the five human dataset folds to each of the five species-specific dataset folds. For human data, we provide five evaluation results obtained on each of the human dataset folds.

### 4.7 Token attribution analysis

We employed the Integrated Gradients algorithm [36] to conduct token attribution analysis. For epigenetic data analysis, we utilized the *bigbird-base-sparse-t2t* model, which was fine-tuned on the standard DeepSEA dataset with sequences of 1,000 bp. Despite the dataset comprising over 900 features, our analysis specifically targeted six key features: ATF1, CTCF, GATA2, H3K27me3, H3K9me3, and H3K4me1 ChIP-seq profiles from untreated K562 cells. For each genomic feature, we randomly chose 3,000 nucleotide sequences that encompassed ChIP-seq peaks. Subsequent tokenization of these sequences adhered to the same methodology as that applied in the chromatin profile fine-tuning task. With default parameters set, token attribution values were derived. For motif analysis, we leveraged the XSTREME tool [64]. Both FIMO and XSTREME assessments sourced motifs from the HOCOMOCO v11 database [65].

For token importance analysis concerning promoter mutations, we utilized the *gena-lm-bert-large-t2t* model fine-tuned on the 2 kb length promoter dataset as previously described. For each mutation, we identified all overlapping promoter regions and calculated token importance scores using Integrated Gradients. For each sample, the top-1 percentile of tokens were designated as ”Highly important”, while the remainder were classified as ”Not important”. We then overlapped mutations with these tokens and categorized each mutation as either ”Overlapping highly important token” or ”Overlapping not important token”. If mutation overlapped tokens from both classes, the ”Highly important token” class was assigned. Finally, we compared the distribution of pathogenic and benign mutations across the ”Overlapping highly important token” and ”Overlapping not important token” classes using a chi-squared test.

### 4.8 Code availability

The code to generate the findings of this manuscript is available in the *supplementary code* section and on our GitHub repository: https://github.com/AIRI-Institute/GENALM. Additionally, our trained models can be found on HuggingFace under the prefix ”gena-lm”: https://huggingface.co/AIRI-Institute/.

## Declarations

### Code availability

The code is available as a ”Supplementary code” file and on GitHub: https://github.com/AIRI-Institute/GENA LM. Pre-trained and fine-tuned models are available with gena-lm-prefix in https://huggingface.co/AIRI-Institute

### Authors’ contributions

M.B., O.K., D.S., V.F., and Y.K. conceived the study; Y.K. performed models pre-training and fine-tuning experiments, including experiments with RMT; V.F. prepared datasets, conducted bioinformatic analysis, and provided biological interpretation of the results; A.S. performed models fine-tuning and benchmarking with help from M.P and Y.K. M.P. and D.P. performed token importance analysis. D.P. analyzed ClinVar data and splice site mutations. N.C. contributed to dataset preparation; M.B. and O.K. supervised the study; all authors contributed to manuscript preparation.

**Supplementary information.** Supplementary Figs. 1–11, Supplementary Tables 1–6, and Supplementary Code file.

## Footnotes

1 https://dnalm.airi.net

2 https://github.com/AIRI-Institute/GENA LM

3 https://huggingface.co/AIRI-Institute

4 For models starting with the gena-lm-prefix, visit: https://huggingface.co/AIRI-Institute/

5 https://ftp.ensembl.org/pub/release-106/

6 https://epd.epfl.ch/EPDnewselect.php

7 https://data.starklab.org/almeida/DeepSTARR/Data/

8 http://deepsea.princeton.edu/media/code/deepsea_train_bundle.v0.9.tar.gz

9 https://github.com/johli/aparent

10 https://github.com/NVIDIA/apex

