## Supplementary Table 1 for "GENA-LM: A Family of Open-Source Foundational DNA Language Models for Long Sequences"

| H3K4me1 | | |
|---|---|---|
| CONSENSUS | EVALUE | SIM_MOTIF |
| DSAGATAAGA | 9.43E-79 | GATA4_HUMAN.H11MO.0.A |
| RVAGATAAVAD | 1.02E-68 | GATA3_HUMAN.H11MO.0.A |
| DRTGAGTCAYH | 8.49E-57 | JUN_HUMAN.H11MO.0.A |
| DVTGAGTCABH | 3.43E-55 | JUNB_HUMAN.H11MO.0.A |
| KRVTGAGTCAYH | 1.01E-52 | FOSL1_HUMAN.H11MO.0.A |
| NDRTGAGTCAYH | 1.71E-49 | FOSL2_HUMAN.H11MO.0.A |
| WGATAARR | 9.40E-48 | GATA1_HUMAN.H11MO.0.A |

| H3K9me3 | | |
|---|---|---|
| CONSENSUS | EVALUE | SIM_MOTIF |
| GGTTTCTCTCCAGTR | 1.10e-078 | ZN274_HUMAN.H11MO.0.A |
| TCCCACATTCA | 2.80e-071 | TBX21_HUMAN.H11MO.0.A |
| AGGCTTTCCCA | 5.20e-027 | NFKB1_HUMAN.H11MO.1.B |
| AGAAYTCACACTGGR | 1.10e-021 | ZNF18_HUMAN.H11MO.0.C |
| YCCCTWGCWGCWAGG | 2.20e-003 | RFX5_HUMAN.H11MO.0.A |
| H3K27me1 | | |
| CONSENSUS | EVALUE | SIM_MOTIF |
| RRCAGGTGCN | 3.88e-013 | SNAI2_HUMAN.H11MO.0.A |
| CHVCASCTGCYBYY | 2.48e-012 | ASCL1_HUMAN.H11MO.0.A |
| SMCAGCTGCWB | 5.64e-010 | TFE2_HUMAN.H11MO.0.A |
| SCAGGTGK | 5.30e-008 | SNAI1_HUMAN.H11MO.0.C |
| BVCAGGTGWG | 2.07e-007 | ZEB1_HUMAN.H11MO.0.A |
| CHSCAGCTGYYYB | 3.09e-007 | MYOG_HUMAN.H11MO.0.B |
| DGCAGSTGKS | 1.51e-005 | ITF2_HUMAN.H11MO.0.C |
