## Supplementary Table 3 for "GENA-LM: A Family of Open-Source Foundational DNA Language Models for Long Sequences"

| Specie | NCBI accession id |
|---|---|
| homo_sapiens | GCF_000001405.40 |
| pan_troglodytes | GCF_028858775.1 |
| mandrillus_leucophaeus | GCF_000951045.1 |
| otolemur_garnettii | GCF_000181295.1 |
| mus_musculus | GCF_000001635.27 |
| cricetulus_griseus | GCF_000223135.1 |
| jaculus_jaculus | GCF_020740685.1 |
| ictidomys_tridecemlineatus | GCF_016881025.1 |
| ursus_americanus | GCF_020975775.1 |
| tursiops_truncatus | GCF_011762595.1 |
| loxodonta_africana | GCF_000001905.1 |
| taeniopygia_guttata | GCF_003957565.2 |
| gallus_gallus | GCF_016699485.2 |
| latimeria_chalumnae | GCF_000225785.1 |
| salmo_salar | GCF_905237065.1 |
| hucho_hucho | GCA_003317085.1 |
| haplochromis_burtoni | GCF_018398535.1 |
| amphiprion_percula | GCA_003047355.2 |
| carassius_auratus | GCF_003368295.1 |
| danio_rerio | GCF_000002035.6 |
| sinocyclocheilus_grahami | GCF_001515645.1 |
| ciona_savignyi | GCA_000149265.1 |
| drosophila_melanogaster | GCF_000001215.4 |
| caenorhabditis_elegans | GCF_000002985.6 |
| saccharomyces_cerevisiae | GCF_000146045.2 |
| arabidopsis_thaliana | GCF_000001735.4 |
| escherichia_coli | GCF_000008865.2 |
