## Supplementary Note for "GENA-LM: A Family of Open-Source Foundational DNA Language Models for Long Sequences"

### Supplementary Fig. 1

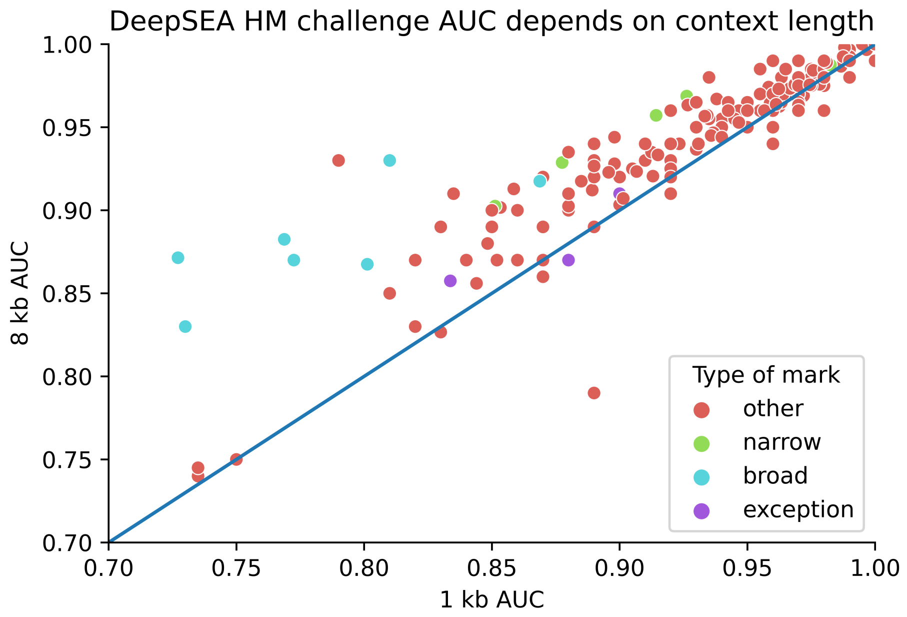


**Supplementary Fig. 1** AUC score comparison for predicting chromatin occupancy, with a breakdown for each specific chromatin mark, contrasting GENA-LM trained on 1-kb versus 8-kb contexts

### Supplementary Fig. 2

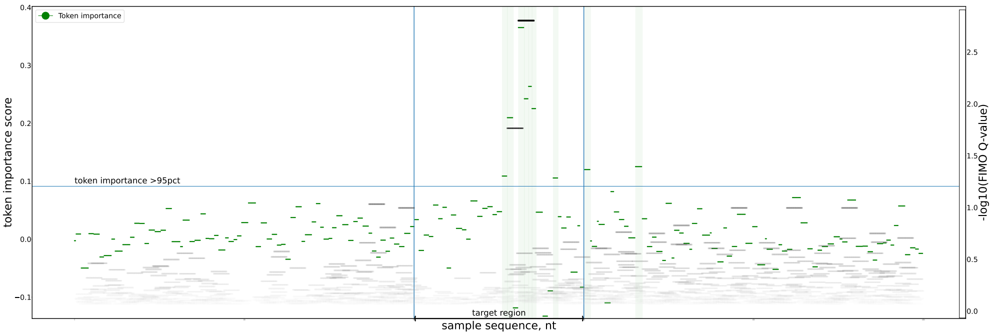


**Supplementary Fig. 2** Distribution of token importance scores and FIMO motif scores for a single representative example of the DeepCT dataset. Horizontal green dashes correspond to tokens, and grey dashes correspond to FIMO motifs. The X-axis shows DNA sequence coordinates (from 0 to 1000 bp). The left Y-axis shows token importance (the higher, the more important). Vertical highlights mark the positions of highly important tokens. The right Y-axis shows FIMO scores (the higher, the more important). FIMO motifs are colored according to their strength (dark gray indicates strong motifs).

### Supplementary Fig. 3

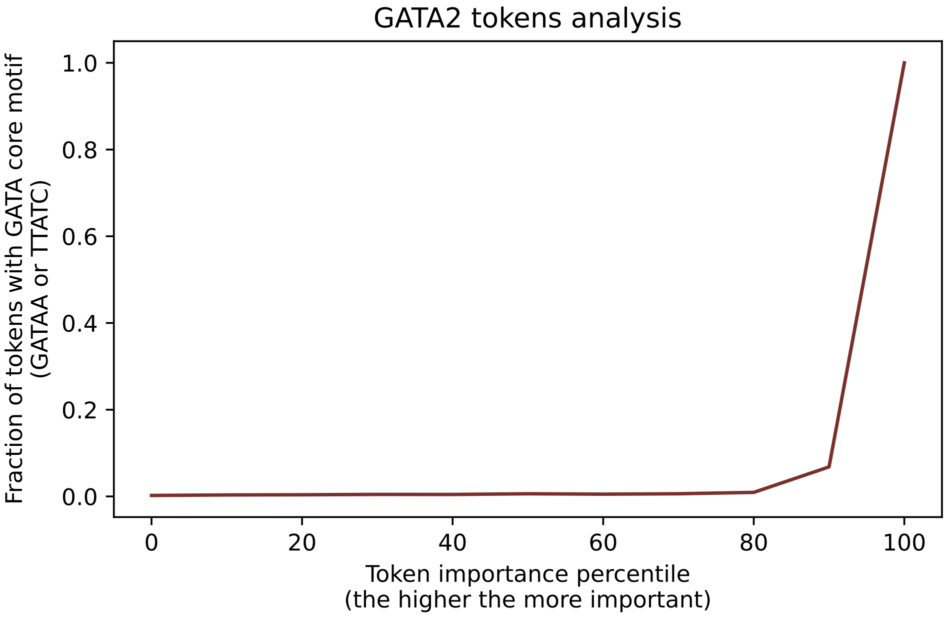


**Supplementary Fig. 3** "GATAA" subsequence is present in tokens highly important for GATA2 binding prediction. The X-axis shows the percentile of token importance, the Y-axis indicates the frequency of "GATAA" subsequence occurrence in the corresponding group of tokens.

### Supplementary Fig. 4

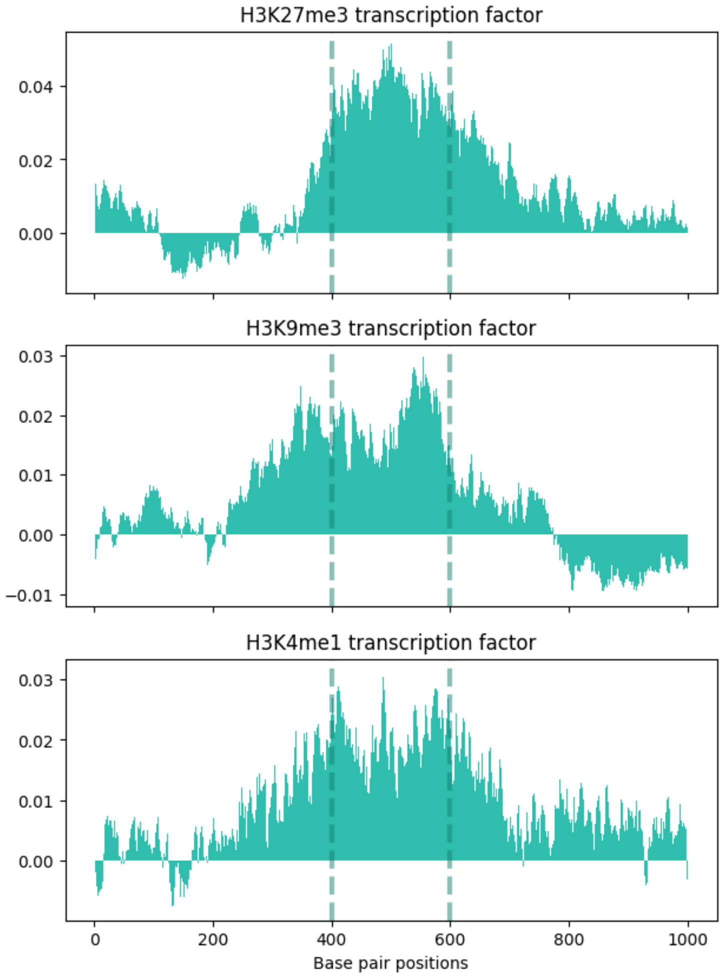


**Supplementary Fig. 4** Distribution of average token importance scores across sequence length. Each row in the panel corresponds to one factor (factor name indicated on the top of plots). The vertical dashed lines indicate the 200 bp region over which the model makes a prediction.

### Supplementary Fig. 5

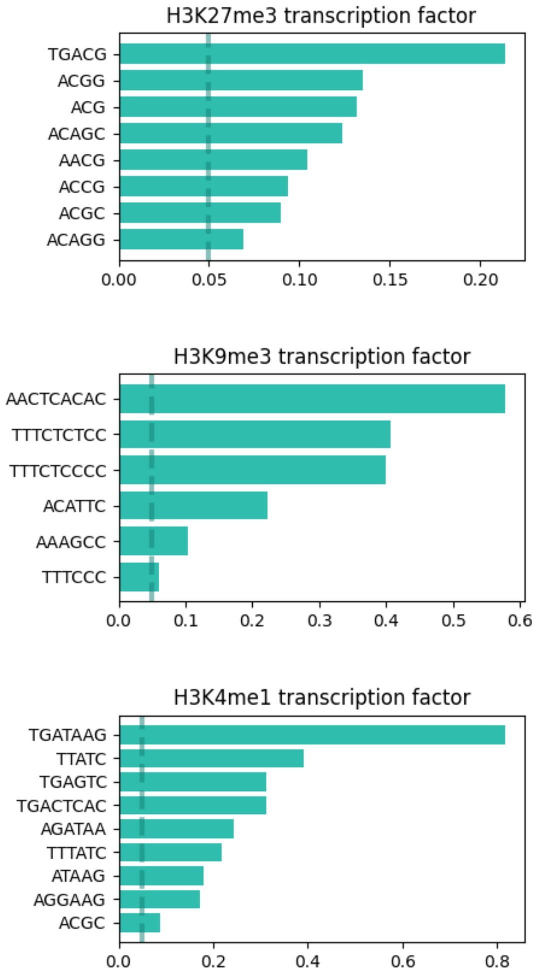


**Supplementary Fig. 5** Bars showing how often a token occurs in the ”highly important” tokens list (list of tokens with importance score in top 5 percentile). The X-axis represents a fraction of token occurrences with a ”highly important” score of all token occurrences. The vertical line indicates a fraction threshold of 0.05. Only tokens characterized by a fraction above this threshold are shown.

#### Supplementary Fig. 6

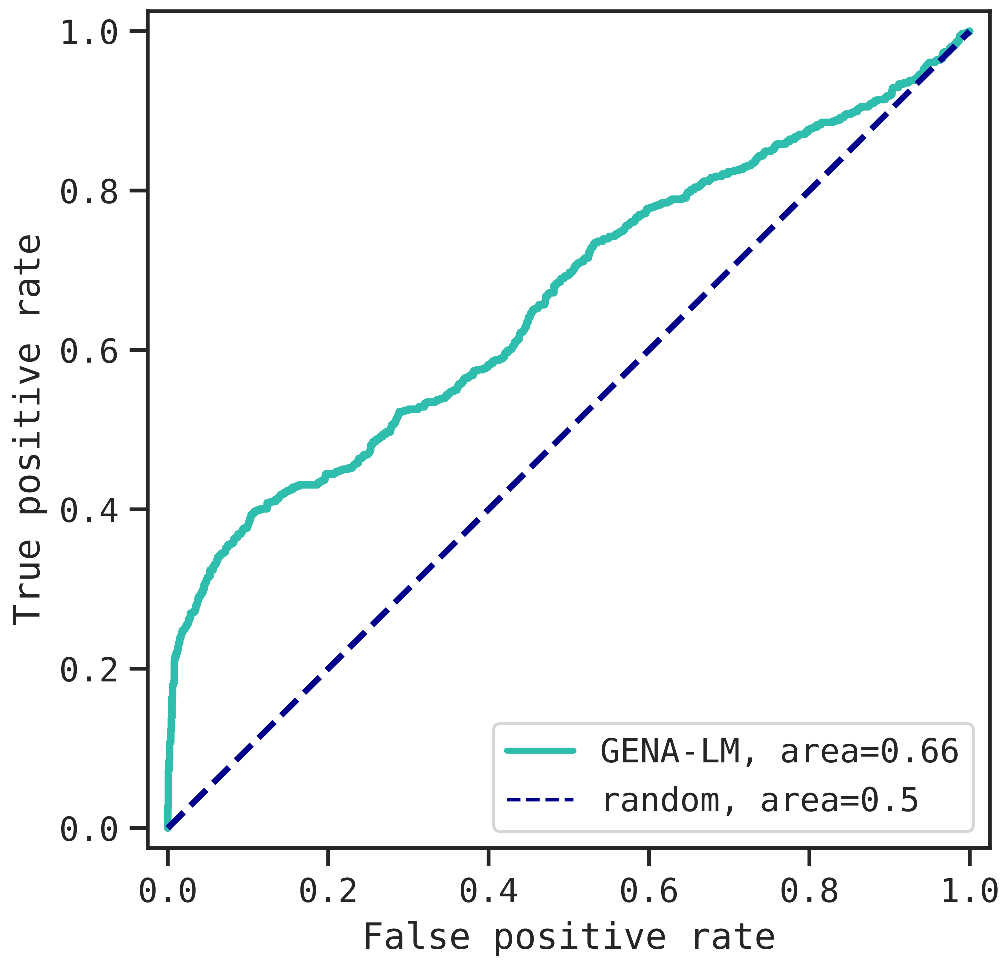


**Supplementary Fig. 6** ROC-AUC curve for zero-shot prediction of the significance of ClinVar regulatory single nucleotide mutations, using the GENA-LM model fine-tuned for the promoter prediction task.

#### Supplementary Fig. 7

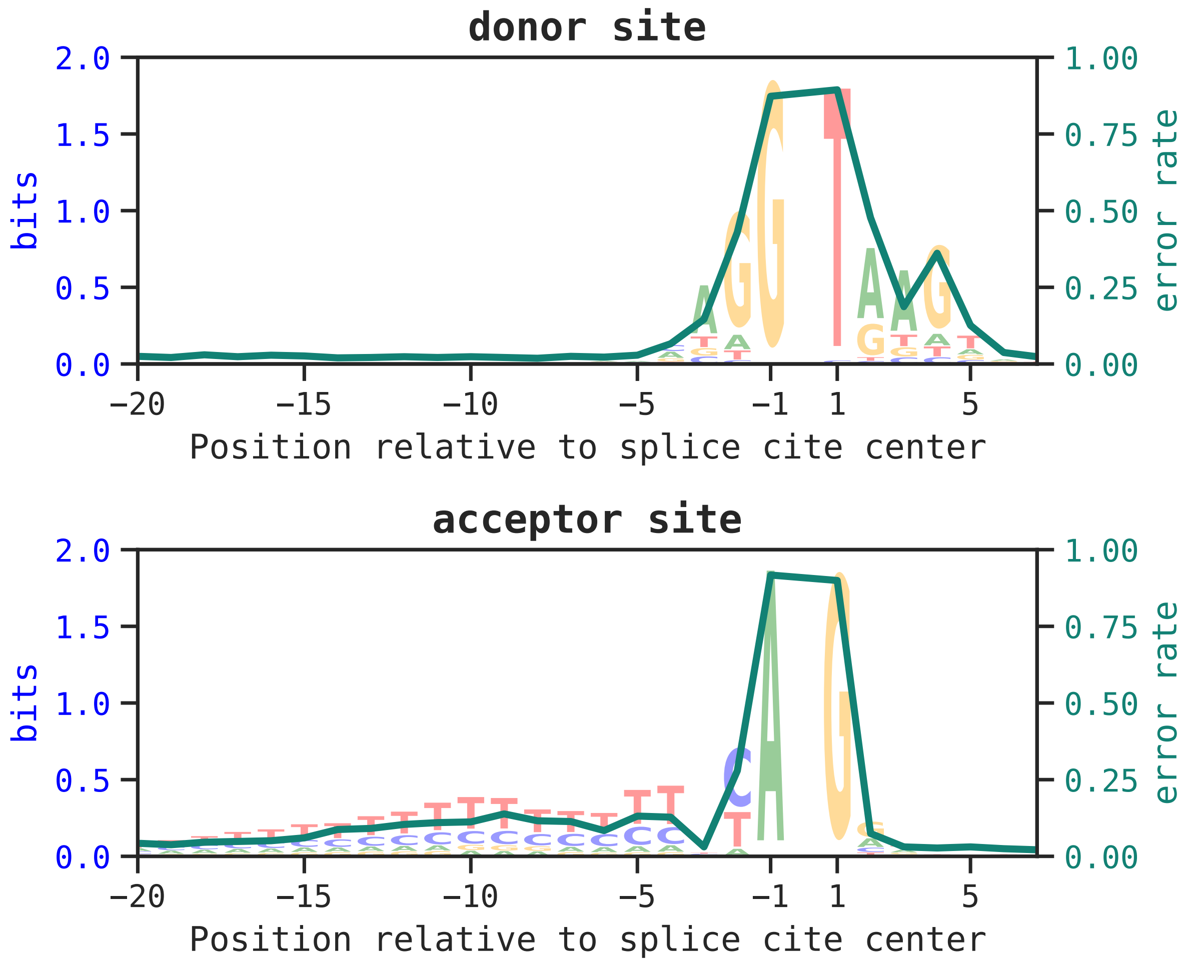


**Supplementary Fig. 7** Correspondence between the effect of a single nucleotide mutation at a position on model prediction and the information content of the position. The left y-axis shows the information content of each position. The right y-axis shows the average number of splice site classification errors due to mutation at position

### Supplementary Fig. 8

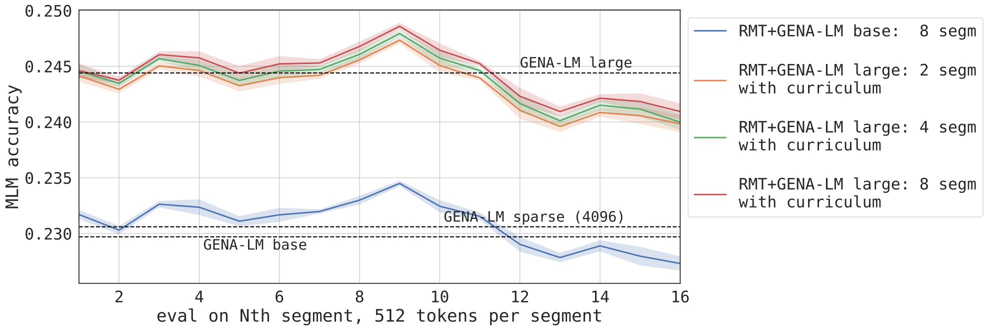


**Supplementary Fig. 8** Effect of RMT application on MLM accuracy. Long DNA sequence input was split into continuous segments, each represented by 512 tokens, and the resulting chain of segments was scanned with RMT from left to right. Information stored in the memory tokens during the processing of the first segments can be used to solve MLM problem in the next segments. The graph shows the dependence of MLM accuracy from segment index. Applying RMT to the base-sized model improves MLM accuracy for 2–10 segments. However, the large model shows no clear benefit from pre-training in terms of MLM accuracy. Similarly, a sparse model, with an input length of 4096 tokens (equivalent to 8 segments), also doesn't show a noticeable advantage from increased context length on MLM accuracy. MLM results are averaged over 5 runs on the valid set with different seeds for masking.

### Supplementary Fig. 9

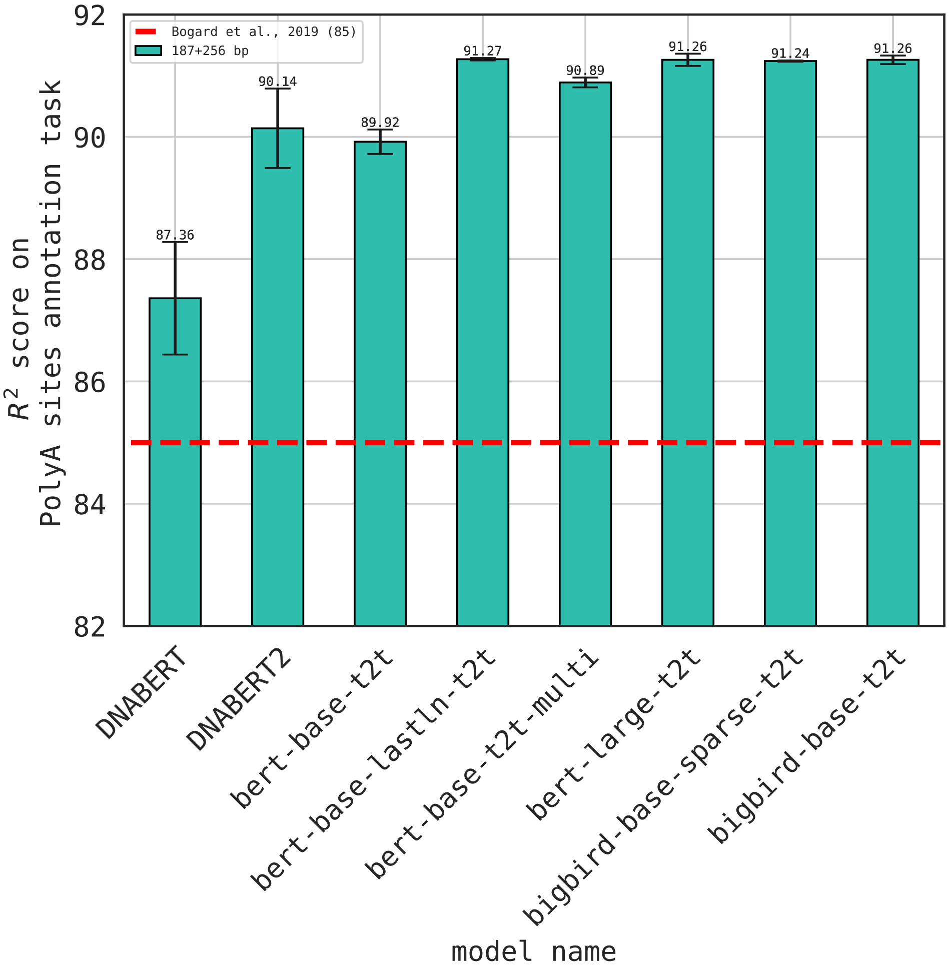


**Supplementary Fig. 9** Benchmarking GENA-LMs on prediction of polyadenylation site strength task

#### Supplementary Fig. 10

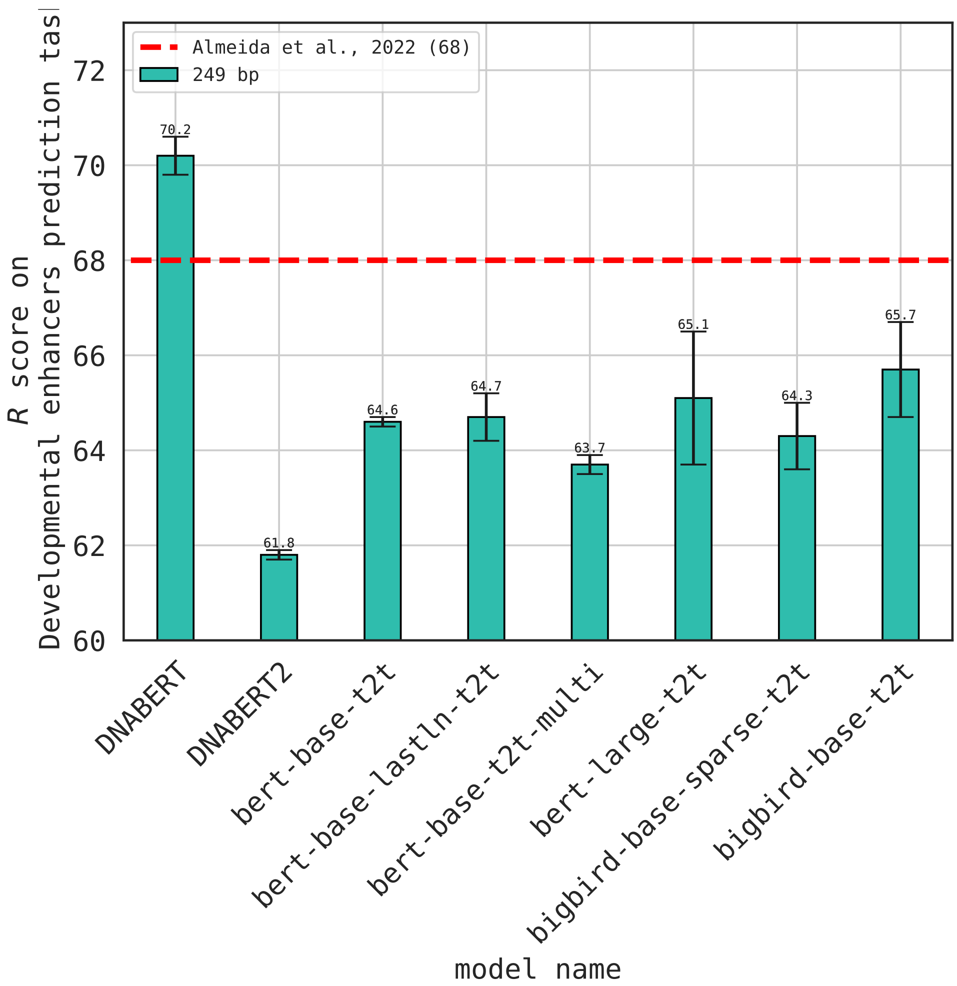


**Supplementary Fig. 10** Drosophila developmental enhancer classification

#### Supplementary Fig. 11

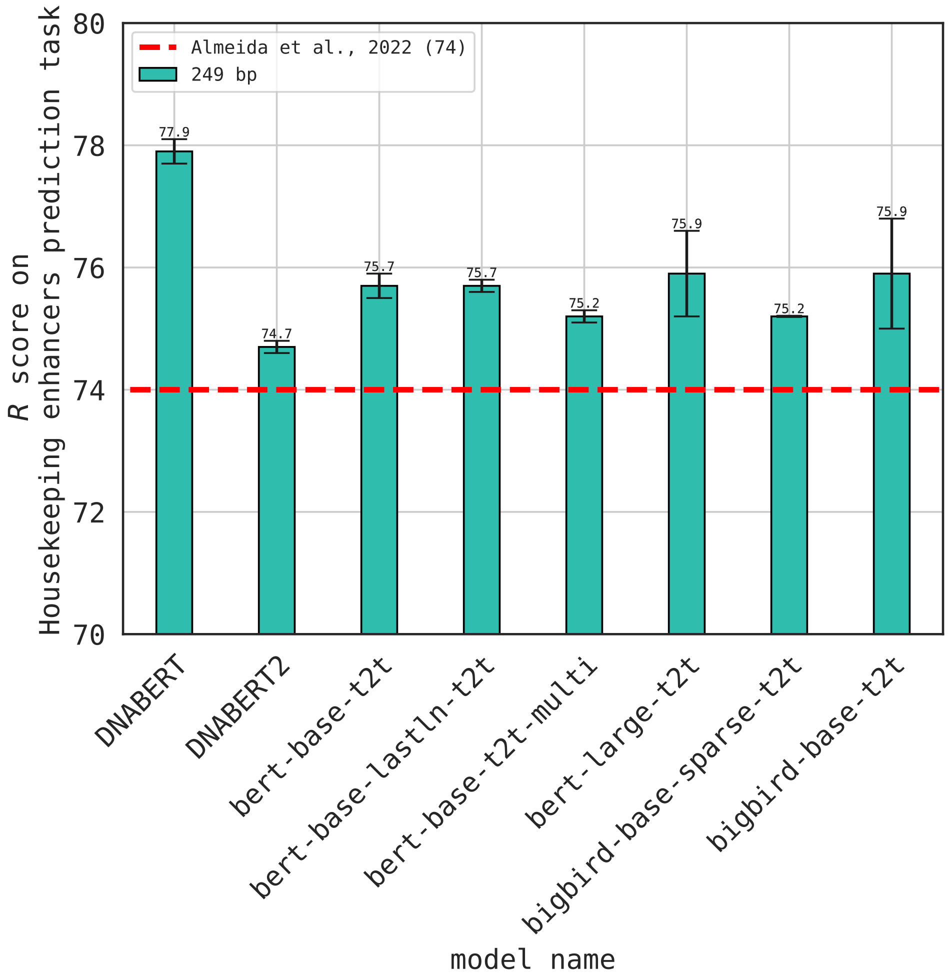


**Supplementary Fig. 11** Drosophila housekeeping enhancer classification

### Supplementary Table 1

Significantly enriched motifs were identified using XSTREME for H3K4me1, H3K9me3 and H3K27me3 histone marks. File `supplementary_table_1.pdf`.

### Supplementary Table 2

The relationship between the importance of a token according to model attributions score (99th quantile) and the clinical significance of single nucleotide mutations in it. File `supplementary_table_2.pdf`.

### Supplementary Table 3

List of species used for phylogenetic analysis with GENA-LMs. File `supplementary_table_3.pdf`.

### Supplementary Table 4

List of species included in the multispecies GENA-LM training setup. File `supplementary_table_4.xlsx`.

### Supplementary Table 5

Detailed information about each model from the GENA-LM family. File `supplementary_table_5.xlsx`.

### Supplementary Table 6

CTCF and H3K27ac dataset sources for the training of interspecies transferability of the human-based model. File `supplementary_table_6.csv`.

### Supplementary Note 1

***Prediction of polyadenylation site strength.***

Polyadenylation in DNA plays a pivotal role, and variants that influence polyadenylation site selection have been linked to diseases. Recently, [1] introduced a reporter assay that quantifies the strength of the proximal polyadenylation signal across millions of short sequences. This study showcased specific nucleotide determinants of the polyadenylation signal strength and demonstrated that this strength could be predicted from the DNA sequence using the convolutional neural network, APARENT. In our work, we adapted GENA-LMs to predict polyadenylation signal strength. As illustrated in Supplemenatary Figure 9, all GENA-LM variants significantly surpassed APARENT, with the top-performing GENA-LM achieving a Pearson $R^2$ value of 0.91 $\pm$ 0.0002 compared to APARENT's $R^2$ = 0.85. The DNABERT and DNABERT-2 models, when fine-tuned on the same dataset, also outperformed APARENT ($R^2$ = 0.87 $\pm$ 0.01 for DNABERT and 0.901 $\pm$ 0.007 for DNABERT-2 versus APARENT's $R^2$ = 0.85). However, its performance was lower than that of the GENA-LMs. These outcomes underscore the capability of the GENA-LM architecture to deliver state-of-the-art performance in specific genomic tasks. Nevertheless, it's worth noting that sequences profiled in the polyadenylation signal strength assay are relatively concise, comprising a 187 bp proximal section and a 256 bp distal part. Such short sequences may not leverage the full potential of long-input GENA-LMs. Consequently, we explored the performance of these models on challenges where the value of extended context could be better evaluated.

***Prediction of Drospila enhancer activity.***

Recently, [2] introduced the DeepSTARR dataset, which presents enhancer activity for millions of short sequences gauged in *Drosophila* cells, classifying each sequence by its housekeeping (cell-type unspecific) and developmental (cell-type specific) enhancer strength. The authors showed that the convolutional neural network DeepSTARR adeptly predicted these activities based on nucleotide sequences. When testing with GENA-LMs, our findings were nuanced (Supplementary Fig. 10 and 11). In the realm of developmental enhancers, the specialized DeepSTARR model surpassed GENA-LMs (optimal GENA-LM Pearson R = 0.657 $\pm$ 0.01 vs. DeepSTARR's 0.68). Yet, for housekeeping enhancers, the tables turned with GENA-LM overshadowing DeepSTARR (optimal GENA-LM Pearson R = 0.768 $\pm$ 0.01 vs. DeepSTARR's 0.74). Crucially, against the benchmarks of the Nucleotide Transformer, GENA-LMs consistently showcased higher scores for both categories (Nucleotide Transformer reported R=0.64 for developmental and R=0.75 for housekeeping). Similarly, GENA-LMs outperform DNABERT-2 in both tasks. Interestingly, DNABERT, a Predecessor of DNABERT-2, shows substantially better results when fine-tuned on the DeepSTARR dataset, providing the scores better than GENA-LMs and origina DeepSTARR CNN for both tasks. A plausible explanation for this phenomenon might be the alignment between the length of input DNA sequences used in this study (249 bp) and the input size of DNABERT (512 bp), which is substantially smaller than the input sizes typically employed during the pretraining phases of GENA-LM, DNABERT-2 or Nucleotide Transformer.
